# Capturing Human Trophoblast Development with Naïve Pluripotent Stem Cells In Vitro

**DOI:** 10.1101/2020.12.17.416800

**Authors:** Shingo Io, Mio Kabata, Yoshiki Iemura, Katsunori Semi, Nobuhiro Morone, Ikuhiro Okamoto, Tomonori Nakamura, Yoji Kojima, Chizuru Iwatani, Hideaki Tsuchiya, Belinda Kaswandy, Eiji Kondoh, Mitinori Saitou, Takuya Yamamoto, Masaki Mandai, Yasuhiro Takashima

## Abstract

Trophoblast are extra-embryonic cells that are essential to maintain pregnancy. Human trophoblasts arise from the morula as trophectoderm (TE), which, after implantation, differentiates into cytotrophoblast (CT), syncytiotrophoblast (ST), and extravillous trophoblast (EVT) composing the placenta. Here we show that naïve, but not primed, human pluripotent stem cells (PSCs) recapitulate trophoblast development. Naïve PSC-derived TE and CT (nCT) recreated the human and monkey TE-to-CT transition. nCT self-renewed as CT stem cells and had the characteristics of proliferating villous CT and CT in the cell column of the first trimester. Notably, although primed PSCs differentiated into trophoblast-like cells (pBAP), pBAP were distinct from nCT and human placenta-derived CT stem cells, exhibiting properties consistent of the amnion. Our findings establish an authentic paradigm for human trophoblast development, demonstrating the invaluable properties of naïve human PSCs. Our system will provide a platform to study the molecular mechanisms underlying trophoblast development and related diseases.

## Introduction

Trophoblasts compose the placenta and are essential extra-embryonic cells from blastocyst through birth. The abnormal differentiation of trophoblasts during early pregnancy is considered to cause placenta-related complications such as pregnancy-related hypertension and intrauterine growth restriction (Burton and Jauniaux, 2017; Malhotra et al., 2016). However, it is difficult to analyse trophoblasts in human uterus, especially during the first few weeks after implantation, due to technical difficulties and ethical restrictions.

Trophoblast development and physiology have been described based on mice, and early lethality (embryonic days (E)9.5 −14.5) in knockout lines is frequently associated with severe placental malformations (Perez-Garcia et al., 2018). However, human and mouse placentation differ in many ways. FGFR2 is expressed in mouse trophectoderm (TE) of blastocyst, while FGFR1, 2, and 3 are not expressed in human TE (Kunath et al., 2014). Trophoblast stem cells (TS cells) cultured in FGF4 were established in mice (Tanaka et al., 1998) and used to investigate trophoblast gene functions (Kuckenberg et al., 2010; Nishioka et al., 2009; Niwa et al., 2005; Perez-Garcia et al., 2018; Senner and Hemberger, 2010), while in humans, TS cells cannot be established by FGF4 (Kunath et al., 2014). Instead, human trophoblast research has relied on cancer cell lines derived from choriocarcinoma and immortalized cell lines, but these lines are unreliable for understanding normal physiology, as they have different differentiation ability, transcriptomes, and other properties compared with human trophoblasts (Ji et al., 2013). In 2018, several groups reported the successful culturing of trophoblasts from 5-9 week human placenta or blastocyst for long periods *in vitro* as two-dimensional TS cells or three-dimensional trophoblast organoids (Haider et al., 2018; Okae et al., 2018; Turco et al., 2018). However, whether these cell lines resemble early-stage trophoblasts during peri-implantation development has not been confirmed.

Another approach to study early trophoblast development is to employ pluripotent stem cells (PSCs). Primed human PSCs have been induced to trophoblast lineages by bone morphogenetic protein 4 (BMP4) treatment (Xu et al., 2002). Subsequently, trophoblast-like cells were generated with cocktails of BMP4, the ALK4/5/7 inhibitor A83-01, and the FGFR inhibitor PD173074 (BAP medium) (Amita et al., 2013; Yang et al., 2015). However, the induced *in vitro* cells expressing T and CDX2 differed in HLA class I and epigenetic properties from *in vivo* trophoblasts (Bernardo et al., 2011). In addition, mouse epiblast stem cells (EpiSC), which are the counterpart of primed human PSCs, do not differentiate into trophoblast lineage (Bernardo et al., 2011). In fact, mouse TS cells were established from naïve mouse PSCs, which correspond to the epiblast (EPI) of blastocysts in preimplantation embryos, by CDX2 over-expression and various other genes or conditions (Kuckenberg et al., 2010; Ng et al., 2008; Niwa et al., 2005).

Recently, naïve human PSCs have been established. These cells have gene expression patterns consistent with human pre-implantation EPI and naïve mouse PSCs, thus representing an earlier stage than conventional primed PSCs (Guo et al., 2016; Stirparo et al., 2018; Takashima et al., 2014; Theunissen et al., 2014). Indeed, two groups have reported that naïve human PSCs can differentiate into TS cells (Cinkornpumin et al., 2020; Dong et al., 2020). However, since naïve PSC-derived TE, which is the starting point for trophoblast development from pre-implantation, was not identified, the process of trophoblast development from the pre-implantation stage and the *in vivo* identity of human TS cells remain unclear.

In the present study, we induced TE from naïve human PSCs to establish an *in vitro* model of the trophoblast lineage from TE. Naïve PSC-derived TE successfully differentiated into cytotrophoblasts, which had similar characters with trophoblasts of cynomolgus monkey *(Macaca fascicularis)* embryos *in vivo*, trophoblasts of cultured human embryos, and placenta-derived TS cells. Furthermore, our analysis unveiled the counterpart of *in vitro* human TS cells as some parts of villous cytotrophoblast (VCT) and cytotrophoblast in cell columns (CCC) of *in vivo* first trimester placenta. Finally, we investigated trophoblast development from TE to cytotrophoblast (CT) during peri-implantation stages using this model.

## Results

### Derivation of trophectoderm-like cells from naïve hPSCs

Human trophoblasts arise from the morula as TE. After implantation, TE differentiates into CT, syiicytiotrophoblast (ST), and extravillous trophoblast (EVT) composing the placenta (Fig. 1A). We investigated whether naïve human PSCs could differentiate into TE (Fig. S1A, S1B). Using previously reported single-cell RNA-sequencing (scRNA-seq) data (Petropoulos et al., 2016; Stirparo et al., 2018), we investigated for cell surface antigens expressed in TE of pre-implantation human blastocyst and selected TACSTD2 and ENPEP as marker genes (Fig. 1B). TACSTD2 is expressed in TE of early (E5) and late blastocyst (E7), while ENPEP expression is increased in TE of late blastocyst. Since signaling transduction is important for the development of embryos, we analysed signaling receptors and inferred the necessary signaling pathways in order to develop an appropriate TE induction medium. We examined Nodal, FGF, JAK/STAT, and BMP signaling receptors, as these are known to be related to early development in mice and humans. The expression of signaling receptors among TE, primitive endoderm (PrE) and EPI were evaluated by FPKM and F values (Fig. S1C). The genes with the highest F value in each signaling pathway were ACVR1B for Nodal signaling, FGFR1 for FGF signaling, IL6R for JAK/STAT signaling, and TGFBR3 for BMP signaling. Next, we measured the expression levels of these receptors during pre-implantation development together with FGFR2, the receptor for FGF4, which is known to be important for the induction of mouse TS cells (Kunath et al., 2014). We found that in TE of human embryo (Stirparo et al., 2018), TGFBR3 was highly expressed, but ACVR1B, FGFR1, FGFR2, and IL6R were not expressed (Fig. S1D). These receptors exhibited similar patterns in cynomolgus monkey TE (Nakamura et al., 2016) (Fig. S1E). However, in mouse TE (Nakamura et al., 2015), Tgfbr3 was not expressed, and Acvr1b, Fgfr1, Fgfr2, and Il6ra were expressed (Fig. S1F). To summarize, the receptor expression was similar between humans and cynomolgus monkeys, but different in mice.

**Fig. 1.**
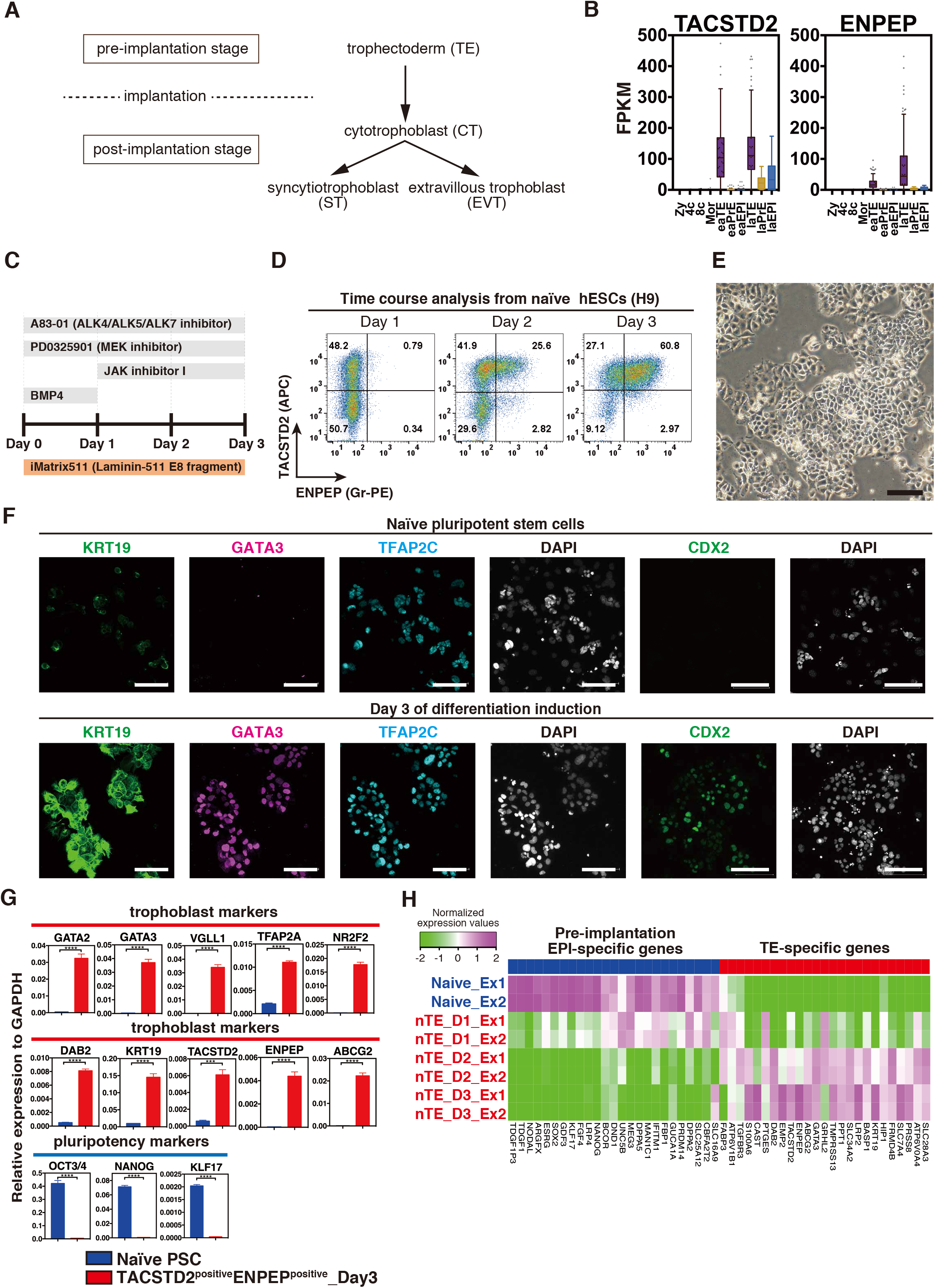
Derivation of trophectoderm-like cells from naïve hPSCs. (A) Schema of trophoblast development. Trophectoderm (TE) in pre-implantation embryos differentiate into cytotrophoblast (CT) after implantation. Post-implantation CT differentiates into syncytiotrophoblast (ST) and extravillous trophoblast (EVT), and composes placenta. (B) Expression in FPKM (fragments per kilobase of transcript per million mapped reads) values of TACSTD2 and ENPEP (CD249) for human early embryo (embryonic days (E)1-7). ENPEP is upregulated in late TE. Zy, zygote; 4c, 4 cells; 8c, 8 cells; Mor, morula; eaTE, early TE (E5); eaPrE, early primitive endoderm ((PrE), E5); eaEPI, early epiblast ((EPI), E5); laTE, late TE (E7); laPrE, late PrE (E7); laEPI, late EPI (E7). Previously reported single-cell RNA-sequencing (scRNA-seq) data was used for the analysis (Stirparo et al., 2018). (C) Schema of the TE induction protocol from naïve human pluripotent stem cells (PSCs) in chemically defined medium. Naïve human PSCs were seeded on Laminin-511 E8 (iMatrix-511) in serum-free basal medium (N2B27) with 2 μM A83-01 (ALK4/ALK5/ALK7 inhibitor), 2 μM PD0325901 (MEK inhibitor), 1 μM JAK inhibitor-1, and 10 ng/ml BMP4. (D) Flow cytometry analysis for the expression of TACSTD2 and ENPEP on days 1-3 after the TE induction of naïve human PSCs. TACSTD2 is expressed on days 1 and 2. ENPEP is expressed on day 3. (E) Bright image of naïve PSC-derived cells on day 3 after the TE induction. Representative images from n=10. Scale bar, 100 μm. (F) Immunofluorescence images of trophoblast markers. Naïve PSCs and naïve PSC-derived cells on day 3 after the TE induction were stained by the trophoblast markers KRT19, GATA3, TFAP2C, and CDX2. Representative images from n=3. Scale bars, 100 μm. (G) Upregulation of trophoblast genes in TACSTD2^+^ENPEP^+^ cells on day 3. Naïve PSC-derived cells on day 3 were sorted by TACSTD2 and ENPEP, and the expression of trophoblast and pluripotency markers was measured by RT-qPCR. (n=3) (H) Relative gene expression dynamics during TE induction. The top 25 EPI- and TE-representative genes were measured by RNA-seq. Human EPI- and TE-representative genes were identified previously (Petropoulos et al., 2016). Naive, naïve PSCs; nTE_D1, and D2, naïve PSC-derived TACSTD2^+^ENPEP^-^ cells on day 1 and day 2, respectively; nTE_D3, TACSTD2^+^ENPEP^+^ cells on day 3; Ex1 or 2, sample1 and 2, respectively.

Based on the above findings, we cultured naïve human PSCs in a cocktail of A83-01, PD0325901 (MEK inhibitor), JAK inhibitor I, and BMP4 for the TE induction (Fig. 1C). In this condition, TACSTD2^positive^ENPEP^negative^ (TACSTD2^+^ENPEP^-^) cells were observed on days 1 and 2, and around 60% of TACSTD2^positive^ENPEP^positive^ (TACSTD2^+^ENPEP^+^) cells were induced on day 3 (Fig. 1D). Cell morphology on day 3 was polygonal and flat, whereas the tightly packed three-dimensional colonies of naïve PSCs were not seen (Fig. 1E, S1A). Immunofluorescence showed that TE marker proteins (KRT19, GATA3, TFAP2C, CDX2) were expressed on day 3 (Fig. 1F) (Blakeley et al., 2015; Chen et al., 2009). We sorted TACSTD2^+^ENPEP^+^ cells on day 3 by flow cytometry and confirmed that TACSTD2^+^ENPEP^+^ cells expressed several trophoblast markers including GATA2, GATA3, VGLL1, TFAP2A, NR2F2, DAB2, KRT19, TACSTD2, ENPEP, and ABCG2, whereas the expression of the pluripotent genes OCT3/4, NANOG, and KLF17 was decreased (Fig. 1G). Further, we performed RNA-seq to analyse global gene expression patterns. The top 25 genes among the EPI-specific and TE-specific genes defined by Petropoulos et al. (Petropoulos et al., 2016) revealed that EPI-specific genes were highly expressed in naïve PSCs, but the expression declined with differentiation (Fig. 1H). Meanwhile, the expression of almost all TE-specific genes was increased on days 2 and 3 (Fig. 1H). The data for TE-specific gene expression suggested that naïve PSCs could differentiate into TE-like cells (nTE).

To verify the generality of our findings, three naïve human ES/iPS cell lines (H9 ESCs, 409B2 iPSCs, and AdiPS iPSCs) underwent two different naïve-induction procedures (chemical reset and 5i/L/A) (Guo et al., 2017; Theunissen et al., 2014). TACSTD2^+^ENPEP^+^ cells were induced in all cases (Fig. S1G).

### Naïve PSC-derived TE differentiates into CT

TE of pre-implantation blastocyst differentiates into CT just after implantation *in vivo.* Therefore, we analysed whether nTE differentiates into post-implantation CT (Fig. 2A). First, we analysed human first trimester chorionic villi (5 to 11 gestational weeks) to search for cell surface antigens and found that TACSTD2 and ENPEP were expressed in CT at 5 gestational weeks (Fig. S2A). SIGLEC6, which has been reported to be expressed in the trophoblast of early pregnancy (Rumer et al., 2012), was detected in CT (CCC and VCT) and ST at 5 gestational weeks (Fig. 2B), but not in TE (Fig. S2B). Flow cytometry confirmed that first trimester placenta (9 and 11 gestational weeks) expressed TACSTD2, ENPEP, and SIGLEC6 (Fig. 2C, S2C), and the placenta-derived TS cell line CT30 (Okae et al., 2018) also expressed all three markers (Fig. S2D). The expression of mouse *Tacstd2, Enpep, Siglec5* (mouse ortholog for human SIGLEC6) was conserved in mouse pre-implantation TE (Fig. S2E). We thus chose to use these three genes as markers for CT. As expected, nTE on day 3 was SIGLEC6^-^ (Fig. 2D).

**Fig. 2.**
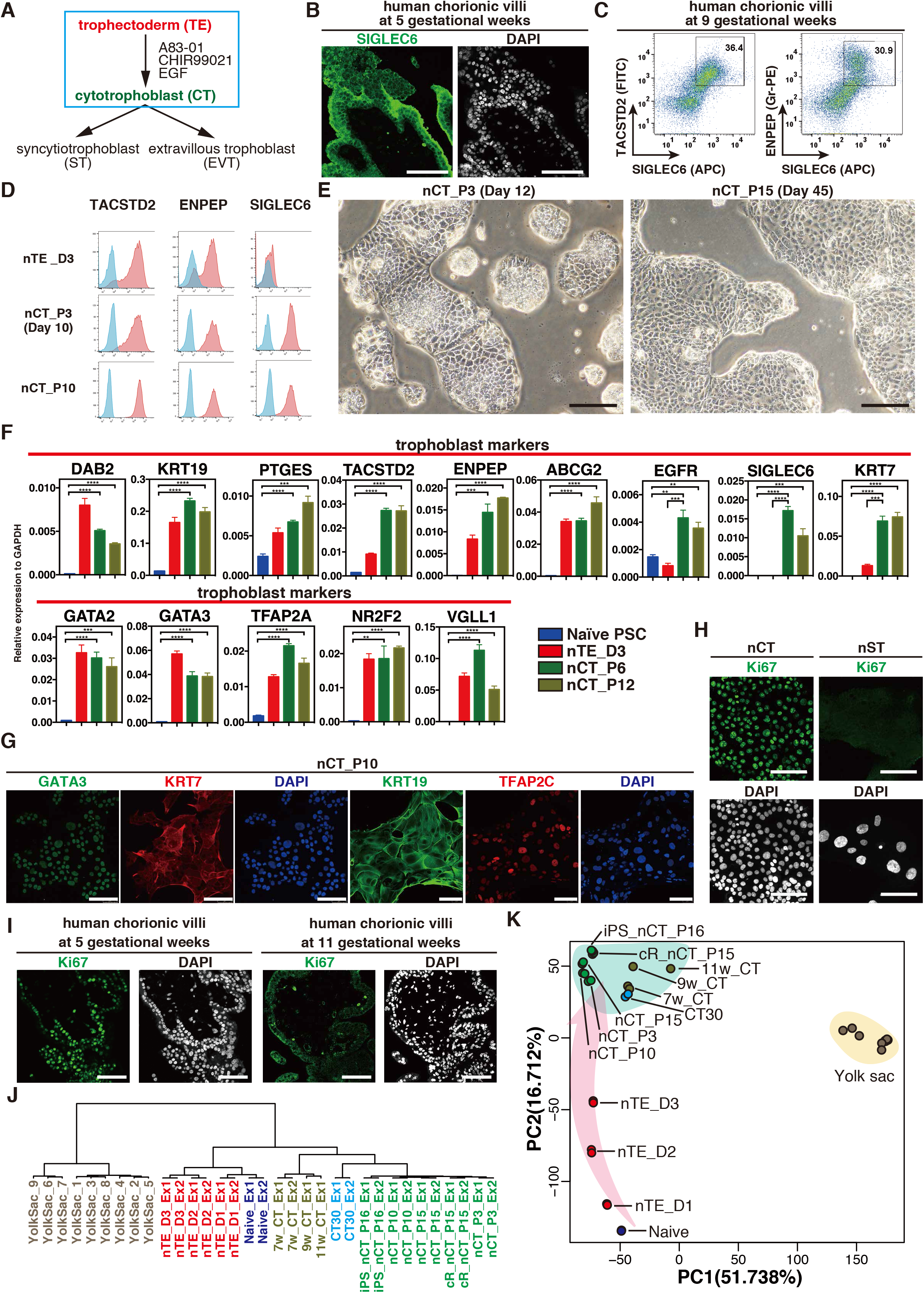
Naïve PSC-derived TE differentiates into CT. (A) Schema of human trophoblast lineages. TE differentiates into CT, which differentiates into both ST and EVT. nTE was differentiated into nCT (nTE-derived CT) with A83-01, CHIR99021, and EGF (ACE) treatment. (B) Immunofluorescence images of human chorionic villi at 5 gestational weeks for SIGLEC6. Scale bars, 100 μm. Representative images from n=3. (C) Flow cytometry analysis for the expression of TACSTD2, ENPEP, and SIGLEC6 on human chorionic villi (9 gestational weeks). (D) Flow cytometry analysis for the expression of TACSTD2, ENPEP, and SIGLEC6 during differentiation from TE to CT. nTE expresses TACSTD2 and ENPEP, but not SIGLEC6. nTE differentiated into nCT in ACE medium on day 10. nCT expresses SIGLEC6. n=6. Similar results were obtained with two independent cell lines. (E) Representative bright images of nCT at passages 3 (left) and 15 (right). Scale bars, 100 μm. Representative images from n=10. (F) RT-qPCR analysis for representative trophoblast markers. *EGFR, SIGLEC6,* and *KRT7* are upregulated strongly only in nCT. The other trophoblast markers are expressed similarly in nTE and nCT. n=3. (G) Immunofluorescence images of nCT for the trophoblast markers GATA3, KRT7, KRT19, and TFAP2C. Scale bars, 100 μm. Representative images from n=3. (H) Immunofluorescence images of nCT and nST for the proliferation marker Ki67. Ki67 was expressed in the nuclei of nCT. Scale bars, 100 μm. Representative images from n=3. (I) Immunofluorescence images of first trimester chorionic villi for the proliferation marker Ki67. Ki67 was expressed in the nuclei of CCC and VCT at 5 gestational weeks. Scale bars, 100 μm. Representative images from n=3. (J) Unbiased hierarchical clustering (UHC) of the transcriptomes. Placental chorionic villi, CT30 and nCT are clustered together. RNA was collected twice in each sample. Naïve, naïve PSCs; nTE_D1 (D2 or D3), one (two or three) day(s) after the induction of naïve PSCs; iPS, 409B2; cR, naïve PSCs established by chemical resetting medium (Guo et al., 2017); 7w_, 9w_, and 11w_CT, TACSTD^+^ENPEP^+^SIGLEC6^+^ placental chorionic villi 7, 9, 11 gestational weeks; CT30, placenta-derived trophoblast stem cells established by Okae et al. (Okae et al., 2018); Yolk Sac, 7-12-gestational-week samples reported by Cindrova-Davies et al. (Cindrova-Davies et al., 2017). Ex1 or 2, independent sample1 and 2, respectively. (K) Principal component analysis (PCA) of the indicated cell types during the TE and CT induction of naïve PSCs. Samples are the same as in Figure 2J.

To determine the induction condition from TE to CT, we observed protein expression and signaling transductions in human chorionic villi (7 gestational weeks) (Fig. S2F). Immunofluorescence confirmed that GATA3 was expressed in CCC, VCT and ST, and that TFAP2A was accumulated in the nuclei of CCC. Previous studies have established human TS cells or trophoblast organoids from placenta (Haider et al., 2018; Okae et al., 2018; Turco et al., 2018). Consistently, we observed that non-phosphorylated β-catenin was localised in the cell membrane of CCC and VCT, suggesting that Wnt is active. Phosphorylated(p)-EGF receptor was strongly expressed in the cell membranes of CCC, VCT and ST, whereas p-SMAD2 was not accumulated in the nuclei of CCC, VCT, or ST (Fig. S2G). We therefore cultured nTE on day 3 in A83-01, CHIR99021, and EGF (ACE) medium (Fig. 2A). In this condition, nTE differentiated into TACSTD2^+^, ENPEP^+^, and SIGLEC6^+^ CT-like cells (nCT) on day 10 (Fig. 2D). TACSTD2^+^ENPEP^+^SIGLEC6^+^ cells were sorted on day 10 and continued to be cultured in ACE medium. nCT continuously expressed TACSTD2, ENPEP, and SIGLEC6 (Fig. 2D), and the morphology of nCT was maintained for more than 10 passages and resembled that of placenta-derived TS cells (Fig. 2E). Their chromosomes were normal (Fig. S2H). We examined gene expression using RT-qPCR and found that nCT on passages 6 and 12 expressed common trophoblast genes and also EGFR, SIGLEC6, and KRT7, which were not expressed in nTE (Fig. 2F). Immunofluorescence showed the expression of the trophoblast markers GATA3, KRT7, KRT19, and TFAP2C in nCT at passage 10 (Fig. 2G). nCT could be maintained for more than 35 passages and over 75 days like CT stem cells (Fig. S2I). In previous reports, Ki67 was expressed in CT (Chang and Parast, 2017; Muhlhauser et al., 1993). Almost all nCT expressed Ki67, suggesting that nCT has high proliferative ability (Fig. 2H). Interestingly, CCC and part of VCT of human chorionic villi expressed Ki67 at 5 gestational weeks, but the number of Ki67 positive cells decreased by 9 gestational weeks and had almost disappeared at 11 gestational weeks, indicating that our nCT correspond to the early stage of CCC and VCT (Fig. 2I, S2J). We also tested each signaling molecule in ACE compounds. Activin A, XAV939, and Gefitinib (EGF receptor tyrosine kinase inhibitor) each reduced the cell growth and viability (Fig. S2K), indicating all three ACE compounds are necessary to maintain nCT.

We also evaluated whether ACE condition could establish TS cells from placenta (Fig. S3A). TACSTD2^+^ENPEP^+^SIGLEC6^+^ cells of placental chorionic villi at 7 and 9 gestational weeks were sorted and cultured in ACE medium (Fig. S3A). The cells were maintained for more than 10 passages in ACE and expressed CT markers (Fig. S3B, S3C).

Next, using RNA-seq, we analysed the global transcriptome of nTE, nCT, TACSTD2^+^ENPEP^+^SIGLEC6^+^ placental chorionic villi at 7, 9 and 11 gestational weeks, and CT30, which we cultured as previously reported (Okae et al., 2018), and also the published RNA-seq data of first trimester human yolk sac (Cindrova-Davies et al., 2017). Unbiased hierarchical clustering (UHC) classified the samples into three groups, yolk sac, nTE and naïve PSCs, and placental chorionic villi, CT30 and nCT (Fig. 2J). Clustering suggests that nCT, CT30 and placental chorionic villi have more similar gene expression profiles comparing to nTE and naïve PSCs which correspond to preimplantation developmental stage. In addition, nCT lines were tightly clustered regardless of the passage number or cell line. Consistently, principal component analysis (PCA) revealed that various nCT lines have similar gene expression patterns to CT30 and chorionic villi (Fig. 2K). PCA further revealed a directional and progressive transition of cellular properties during nCT induction (Fig. 2K). nTE was distant from nCT and placental chorionic villi.

We cultured nCT in suspension using a recently reported method for trophoblast organoids (Turco et al., 2018). In this condition, the morphology of nCT-derived organoids (nCT-organoids) was extremely similar to that of trophoblast organoids derived from placenta (Fig. S4A). nCT-organoids were maintained for 2 months and more than 10 passages. RT-qPCR revealed that both CT markers and ST markers were expressed, as previously reported (Haider et al., 2018; Turco et al., 2018) (Fig. S4B). Immunofluorescence demonstrated that nCT-organoids expressed trophoblast markers including KRT7, GATA3, and TFAP2A (Fig. S4C). It was reported that the outer layer of trophoblast organoids was composed of CT-like cells (Haider et al., 2018; Turco et al., 2018). Indeed, the surface of nCT-organoids expressed ITGA6, which is a marker of CT (Okae et al., 2018) (Fig. S4C). We thus revealed that nCT could be maintained the same way as placenta-derived three-dimensional organoids.

Because our highly proliferated nCT had similar gene expression and morphology as CT30 and three-dimensional trophoblast organoids, we concluded that nCT are nTE-derived CT-like stem cells and the counterpart of proliferative CCC and VCT of human chorionic villi at early gestation.

### nTE-derived CT differentiates into ST and EVT

CT is the bipotential progenitor of ST and EVT *in vivo*. We checked the differentiation capacity of nCT into ST and EVT using a previously reported induction medium (Okae et al., 2018) (Fig. 3A). In Forskolin medium, nCT differentiated into multinucleated syncytia with a tentacular structure on the surface (Fig. 3B). This morphology is similar to ST. The fusion index revealed that more than 80% of the cells were multinucleated nTE-derived ST-like cells (nST) after the induction (Fig. 3C). Scanning electron microscopy showed that nST had similar microvilli as *in vivo* ST (Benirschke et al., 2012) (Fig. 3D). RT-qPCR showed that several ST genes including CGA, CGB, PSG3, and SDC1 were upregulated in nST (Jokimaa et al., 1998) (Fig. 3E), and immunofluorescence showed that hCGβ, SDC1, and hPL were expressed (Fig. 3F). In contrast, Ki67, which was abundantly expressed in nCT, was not expressed in nST, which is consistent with ST at 5 gestational weeks, suggesting that nST lost its proliferative ability (Fig. 2H, 2I).

**Fig. 3.**
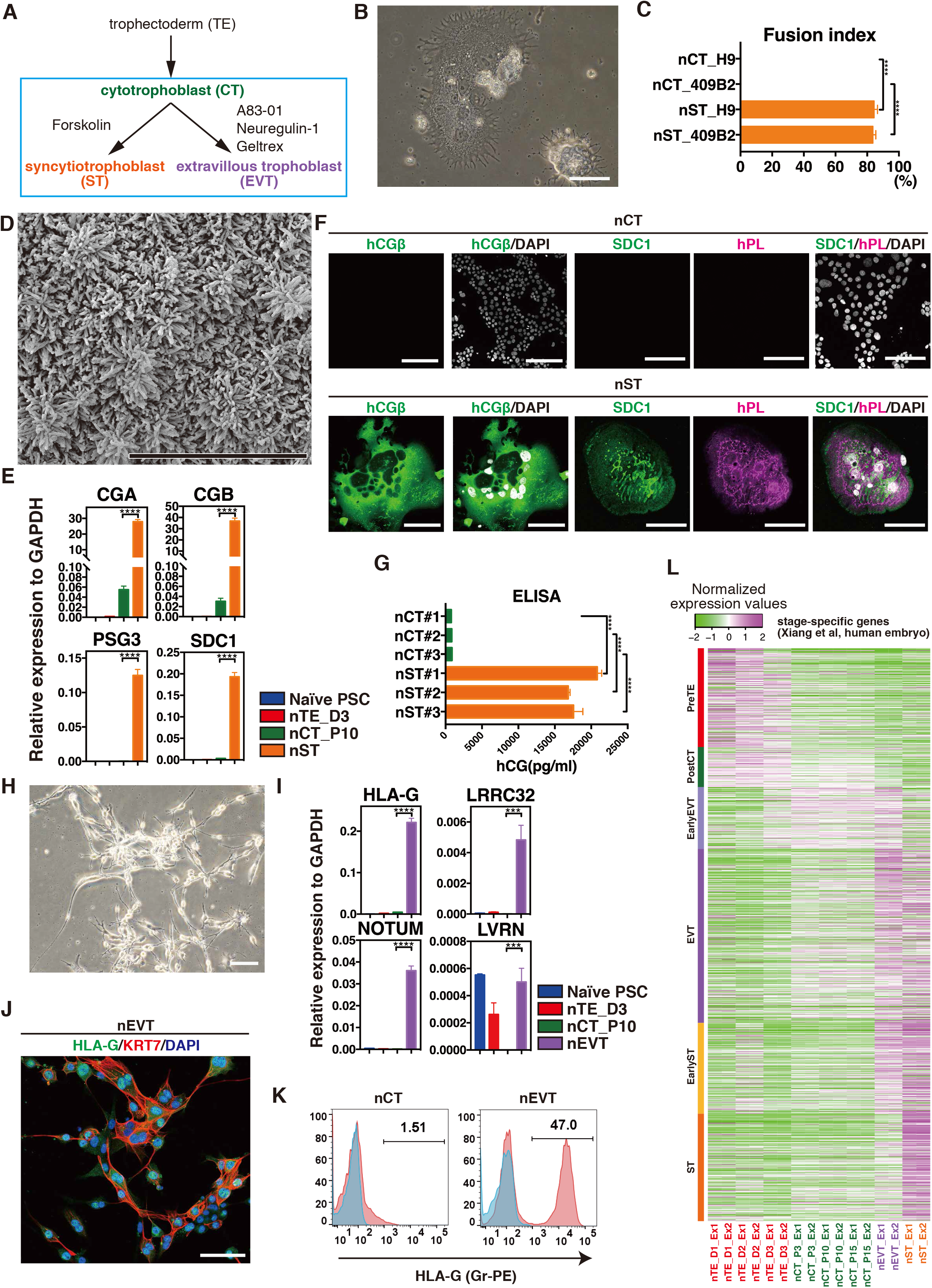
nCT differentiates into ST and EVT. (A) Schema of ST and EVT induction. nCT was maintained in ACE medium. For ST induction, nCT was cultured in Forskolin for 6 days. For EVT induction, nCT was cultured in A83-01, Neuregulin-1, and Geltrex for 8 days. (B) Bright image of nST. nCT was differentiated into ST-like cells (nST) in Forskolin. Representative image from n=10. Scale bar, 100 μm. (C) Fusion efficiency of nST. The fusion index was calculated by the following formula [(number of nuclei in syncytia - number of syncytia) / total number of nuclei x 100]. nST derived from H9 ES cells and 409B2 iPS cells were analysed. Data are presented as the mean + SD (n=3). (D) Scanning electron microscopy image of nST. Microvilli cover the nST surface. Scale bar, 10 μm. Representative images from n=3. (E) RT-qPCR analysis for *CGA, CGB, PSG3,* and *SDC1.* These ST marker genes were upregulated in nST. n=3. (F) Immunofluorescence images of nCT and nST for hCG, SDC1 and hPL. nST are multinuclear, and nuclei are shown in white. Scale bars, 100 μm. Representative images from n=3. (G) Secretion of hCG from nCT and nST. hCG concentration was measured by ELISA. Data are presented as the mean + SD. n=3. (H) Bright image of nEVT. nCT was differentiated into nEVT in A83-01, Neuregulin-1, and Geltrex. Scale bar, 100 μm. Representative image from n=10. (I) RT-qPCR analysis for *HLA-G, LRRC32, NOTUM,* and *LVRN.* These EVT marker genes are upregulated in nEVT. n=3. (J) Immunostaining of nEVT for HLA-G and KRT7. Scale bar, 100 μm. Representative image from n=3. (K) Flow cytometry analysis of the expression of HLA-G in nCT and nEVT. n=3. (L) Relative gene expression of stage-specific genes during peri-implantation. Single cell-RNAseq data of cultured human embryos defined PreTE-, PostCT-, Early EVT-, EVT- and Early ST-, ST-specific genes during the peri-implantation stage (Xiang et al, 2020). Heat maps revealed that nTE, nCT, nEVT, and nST upregulated respective cell-type-specific genes. PreTE, pre-implantation trophectoderm; PostCT, post-implantation CT.

Zhou et al. cultured human embryos *in vitro* from pre-implantation E6 until E14 of post-implantation and showed that trophoblasts on E12 and E14 contained CT and ST (Zhou et al., 2019). Using their defined specific genes for CT and ST on E12 and E14, we analysed the relative gene expression between nCT and nST and found that nCT up-regulated CT-specific genes and nST up-regulated ST-specific genes (Fig. S5A). Finally, we found hCG was secreted from nST into the culture medium (Fig. 3G).

We also cultured nCT in EVT medium containing A83-01 and Neuregulin-1 with Geltrex (Fig. 3A), yielding EVT-like cells (nEVT). The morphology of nEVT was spindle-shape (Fig. 3H). We found that nEVT expressed HLA-G, LRRC32, NOTUM, and LVRN, which were reported to be expressed in EVT of human placenta (Fig. 3I). HLA-G is a marker of EVT. The expression of HLA-G in nEVT was confirmed by flow cytometry and immunostaining (Fig. 3J, 3K). KRT7 was also expressed in nEVT according to immunofluorescence (Fig. 3K).

Next we compared the gene expression patterns of nTE, nCT, nST, and nEVT to *in vitro* cultured human embryos beyond implantation (Xiang et al., 2020). A heat map of gene expression indicated that nTE expressed pre-TE-specific genes, nCT downregulated pre-TE-specific genes and expressed post-CT-specific genes, nEVT upregulated EVT-specific genes, and nST expressed ST-specific genes (Fig. 3L). Furthermore, we compared our cells to first-trimester CT, ST, and EVT (Vento-Tormo et al., 2018). We assigned high-confidence stage marker genes by the expression and distribution in the scRNA-seq data of the placenta and observed the expression of marker genes. We found nCT expressed CT-marker genes. After differentiation, nST and nEVT lost the expression of CT genes and increased the expression of ST and EVT genes, respectively (Fig. S5B).

These data showed that nTE could differentiate into CT lineage that had similar gene expression as post-implantation CT and first-trimester CT. We also observed that nCT differentiates into ST and EVT in differentiation medium even after more than 30 passages (Fig. S5C, S5D). We thus succeeded in establishing a differentiation model to observe trophoblast development from pre-implantation to post-implantation *in vitro*.

### Primed PSC-derived TACSTD2^+^ENPEP^+^ cells do not satisfy trophoblast criteria

Consistent with previous reports (Nakamura et al., 2016), primed human PSCs corresponded to post-implantation EPI (Fig. S6A, S6B). Primed human PSCs have been reported to differentiate into trophoblast-like cells in BAP medium, which contains BMP4, A83-01, and PD173074 (Amita et al., 2013; Yang et al., 2015) (Fig. 4A). Indeed, in BAP medium, primed PSCs differentiated into cells that expressed TACSTD2 and ENPEP on days 2 and 3 (Fig. 4B, 4C).

**Fig. 4.**
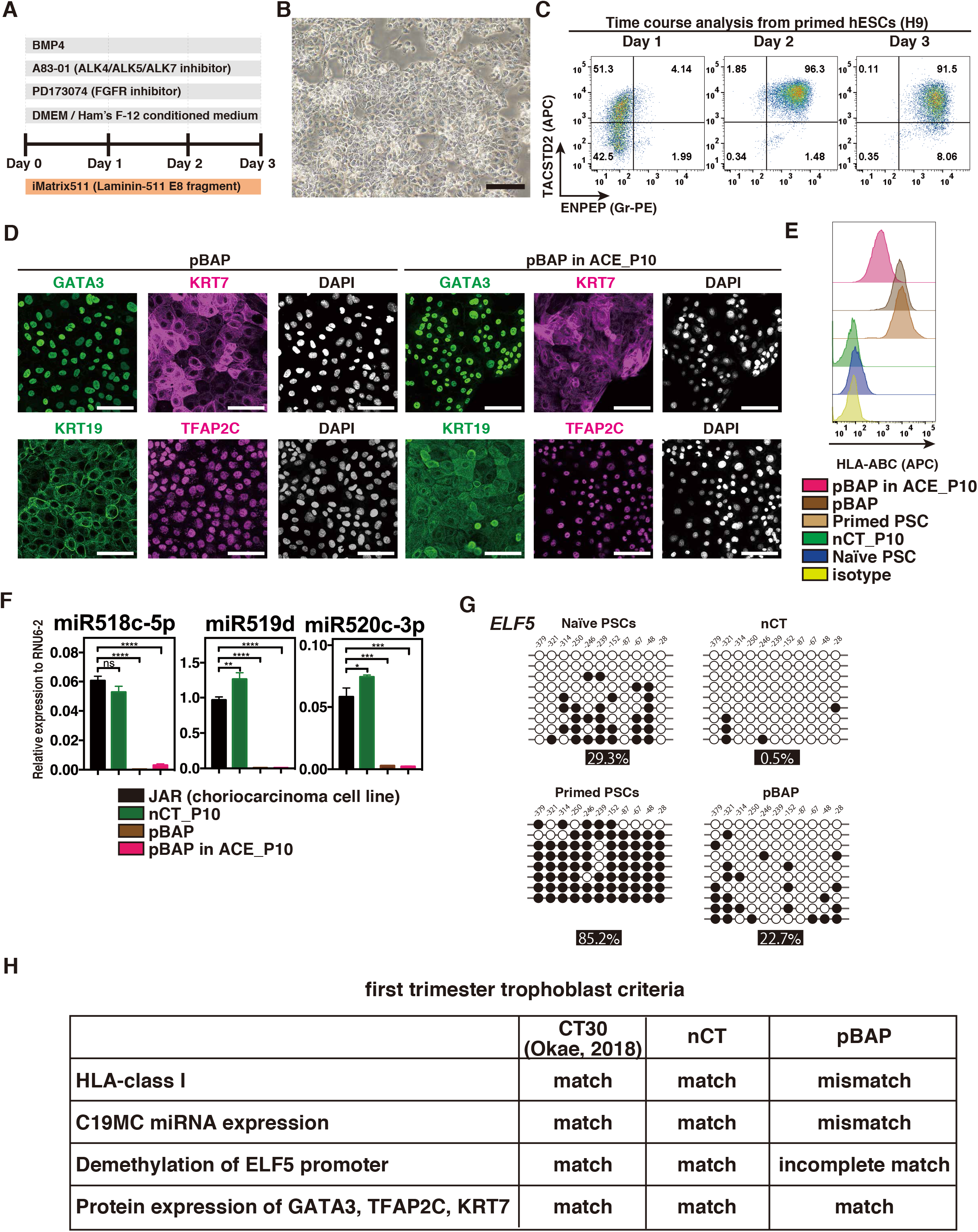
Primed PSC-derived TACSTD2^+^ENPEP^+^ cells do not satisfy trophoblast criteria. (A) Schema of the pBAP induction protocol from primed human PSCs. Primed human PSCs were cultured on Laminin-511 E8 (iMatrix-511) in MEF-conditioned medium with BAP: 10 ng/ml <u>B</u>MP4, 1 μM <u>A</u>83-01 (ALK4/ALK5/ALK7 inhibitor), and 0.1 μM <u>P</u>D173074 (FGFR inhibitor) for 3 days. (B) Bright image of BAP-treated primed PSC-derived cells (pBAP) on day 3. Scale bar, 100 μm. Representative images from n=10. (C) Flow cytometry analysis of the expression of TACSTD2 and ENPEP on days 1-3. TACSTD2 is expressed on day 1. TACSTD2 and ENPEP are expressed on days 2 and 3. Representative images from n=3. (D) Immunofluorescence images of pBAP on day 3 after the induction of primed PSCs and pBAP in ACE medium (passage 10). Primed PSCs were cultured in BAP for 3 days and then cultured in ACE for 10 passages. GATA3, KRT7, KRT19, and TFAP2C are shown. Scale bars, 100 μm. Representative images from n=3. (E) Flow cytometry analysis of the expression of HLA-ABC in naïve PSCs, nCT passage 10, primed PSCs, pBAP, and pBAP in ACE medium (passage 10). n=3. (F) The expression of chromosome 19 microRNA (miRNA) cluster (C19MC) miRNA in JAR (choriocarcinoma cell line), nCT (passage 10), pBAP on day 3, and pBAP in ACE medium (passage 10). The expression was measured using RT-qPCR. n=3. (G) Methylation status of individual CpG sites at the *ELF5* promoter in naïve PSCs, nCT, primed PSCs, and pBAP. Bisulfite sequencing of the *ELF5* promoter region was performed. Open circles show non-methylated CpG and closed circles show methylated CpG. Percentage values below indicate the ratio of methylation (at least 8 clones for each CpG). nCT (passage 10) was demethylated completely, but pBAP on day 3 was demethylated only partially. (H) Criteria for human first trimester trophoblast. CT30 and nCT meet all criteria, but pBAP on day 3 does not. The data of CT30 were published in (Okae et al., 2018).

However, BAP-treated primed PSCs (pBAP) differ from primary trophoblast cells in certain respects (Bernardo et al., 2011). Lee et al. proposed four criteria for human first trimester trophoblast based on the observation of primary trophoblast cells (Lee et al., 2016). We used these criteria of trophoblast to evaluate nCT that we induced from nTE and pBAP that we induced from primed PSCs. pBAP expressed GATA3, KRT7, KRT19, and TFAP2C (Fig. 4D) and could be cultured in ACE medium for 10 passages (Fig. S6C). Even after 10 passages, pBAP in ACE medium continuously expressed these genes, similar to pBAP cultured in BAP medium and nCT (Fig. 2G, 4D). However, flow cytometry analysis for HLA class I revealed that nCT was negative, while pBAP was positive (Fig. 4E). Regarding the expression of chromosome 19 miRNA cluster (C19MC) miRNA, nCT showed a similar level of expression as the choriocarcinoma cell line JAR, but the expression was significantly lower in pBAP (Fig. 4F). Further, nCT showed complete demethylation of the *ELF5* promoter region (0.5%), while pBAP showed incomplete demethylation (22.7%) (Fig. 4G). Even after 10 passages, pBAP in ACE medium showed the same expression patterns of HLA class I and C19MC as pBAP cultured in BAP medium (Fig. 4E, 4F).

From the above, we concluded that nCT satisfied the first trimester trophoblast criteria as CT30 (Okae et al., 2018), but pBAP did not (Fig. 4H).

### nCT represents post-implantation CT of cynomolgus monkey and *in vivo* humans

In order to understand more detailed features of nTE, nCT, and pBAP, we first evaluated the correlation with pre-implantation human embryos by RNA-seq. As previously reported (Nakamura et al., 2016; Stirparo et al., 2018; Takashima et al., 2014), naïve PSCs had high correlation with EPI (Fig. 5A). nTE had particularly high correlation with human embryonic TE (0.82, Figure 5A). The correlation for pBAP with embryonic TE was lower (0.68, Fig. 5A). We also analysed recently reported expanded potential stem cells (EPSCs) and EPSC-derived cells (Gao et al., 2019). However, the correlation for these cells was the lowest (0.56) in three culture conditions (Fig. 5A).

**Fig. 5.**
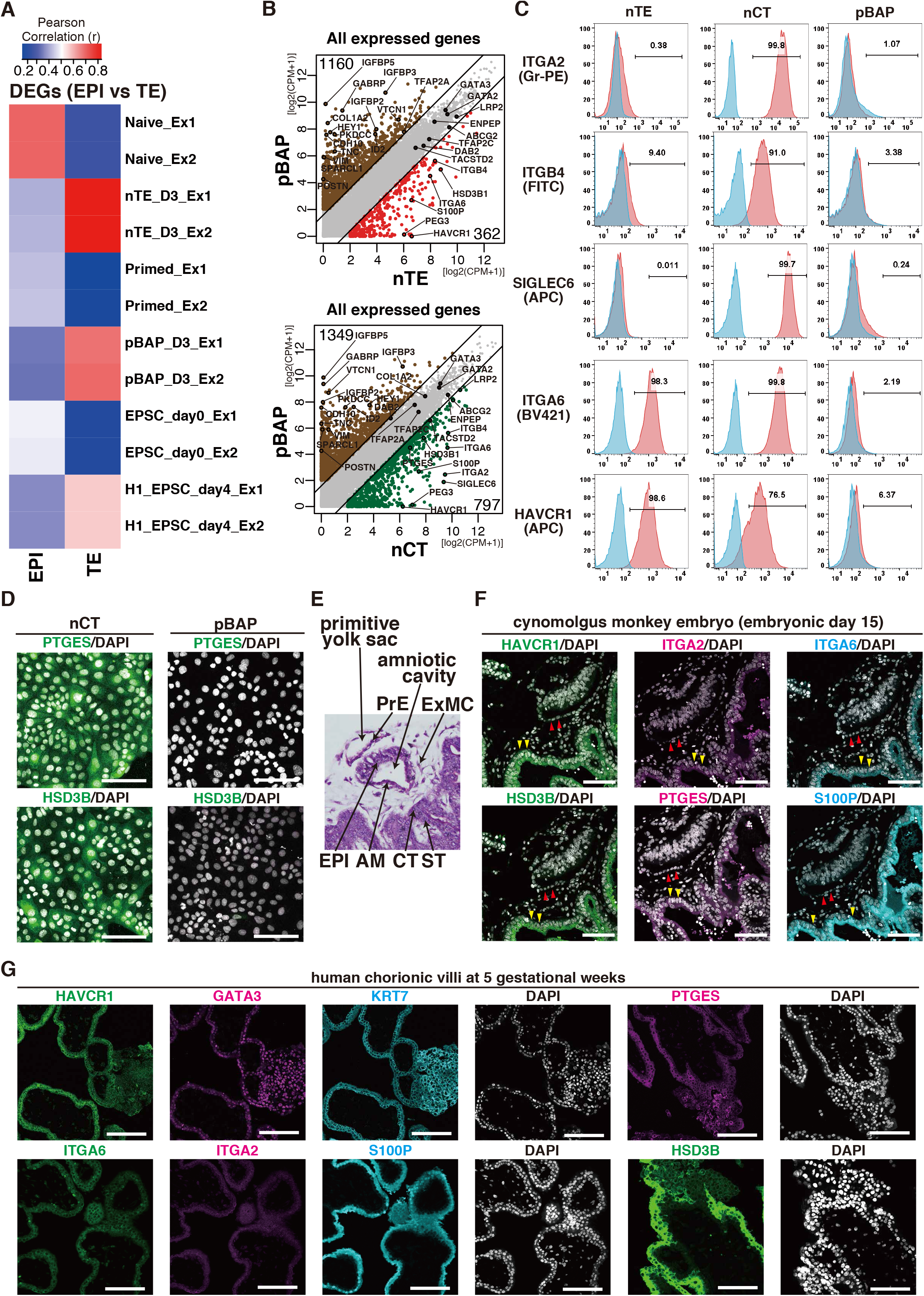
nCT represents post-implantation CT of *in vivo* humans and cynomolgus monkeys. (A) Correlation coefficients of human EPI and TE in pre-implantation embryos with naïve PSCs, nTE, primed PSCs, pBAP, and expanded potential stem cells (EPSCs) (days 0 and 4). Analysis was performed on the differentially expressed genes (DEGs) of EPI and TE in human embryos previously reported (Stirparo et al., 2018). (B) Scatter plots of the average gene-expression levels between nTE (day 3) and pBAP (day 3) (top) and between nCT (passage 3) and pBAP (day 3) (bottom). Brown circles indicate enriched genes in pBAP, red circles indicate enriched genes in nTE, and green circles indicate enriched genes in nCT. Solid black lines indicate 3-fold changes. Key genes are annotated, and the number of enriched genes are indicated. CPM: counts per million. (C) Flow cytometry analysis of nTE, nCT, and pBAP for the key trophoblast-enriched genes identified in Figure 5B. Blue shows an unstained control. Representative images from n=3. (D) The protein expression of PTGES and HSD3B in nCT and pBAP. PTGES and HSD3B were identified as the key trophoblast-enriched genes in Figure 5B. Scale bars, 100 μm. (E) Hematoxylin-Eosin staining of paraffin sections of cynomolgus embryos on embryonic day 15 (E15). AM; amniotic cell, CT; cytotrophoblast, EPI: post-implantation epiblast, ExMC: extraembryonic mesenchyme, PrE; post-implantation primitive endoderm, ST; primitive syncytiotrophoblast. (F) Immunofluorescence images of paraffin sections of cynomolgus embryos on embryonic day 15 (E15). HAVCR1, ITGA2, ITGA6, HSD3B, PTGES, and S100P, which are shown in Figure 5B-E, are expressed in CT and ST of E15 cynomolgus monkey *in vivo.* Scale bars, 100 μm. Yellow arrowheads indicate trophoblast. Red arrowheads indicate amnion. (G) Immunofluorescence images of human chorionic villi at 5 gestational weeks. HAVCR1, GATA3, KRT7, ITGA6, ITGA2, S100P, PTGES, and HSD3B are shown. Scale bars, 100 μm. Representative images from n=3.

Next, we compared the gene-expression levels between nTE and pBAP and between nCT and pBAP by scatter plots (Fig. 5B). GATA2, GATA3, TFAP2C, TACSTD2, and ENPEP were commonly expressed in nTE, nCT, and pBAP (Fig. 5B). Conversely, we found that 362 genes in nTE, 797 genes in nCT, and over 1000 genes in pBAP were highly expressed (Fig. 5B). Among the enriched genes of nCT was SIGLEC6, which we used as a marker of nCT, ITGA6, which is widely used as a CT marker (Okae et al., 2018), and ITGA2, which was recently reported as being expressed in proliferating trophoblasts at the base of CCC (Lee et al., 2018). Furthermore, we identified that HAVCR1, HSD3B1, S100P, PEG3, PTGES, and ITGA6 were upregulated in both nTE and nCT, and ITGA2 and SIGLEC6 were only expressed in nCT. These expression patterns were confirmed by RT-qPCR (Fig. S6D).

We also examined protein expression in nTE, nCT and pBAP. Flow cytometry analysis showed that ITGA2, ITGB4, and SIGLEC6 were expressed only in nCT, and that ITGA6 and HAVCR1 were expressed both in nTE and nCT but not in pBAP (Fig. 5C). Immunofluorescence revealed that PTGES and HSD3B were expressed in nCT but not in pBAP (Fig. 5D). These results confirmed that nCT and pBAP are different cell types.

Next we asked if cells corresponding to nCT exist during early post-implantation *in vivo.* Since we cannot study early post-implantation CT in human uterus, we analysed the embryos of non-human primate cynomolgus monkeys. We isolated early post-implantation cynomolgus monkey embryos (E15) as described previously with ethical approval (Nakamura et al., 2016). Implanted embryos had a bilaminar disc-like structure with amniotic cells and rapidly growing CT and ST (Fig. 5E). We examined the expression of nCT-enriched genes by immunofluorescence. We found HAVCR1, ITGA2, ITGA6, HSD3B, PTGES, and S100P were expressed in CT of cynomolgus monkeys at E15 (Fig. 5F).

To compare nCT to CT in human, we analysed human chorionic villi at 5 gestational weeks, which is a later stage than our observations of cynomolgus monkey. GATA3 and KRT7 marked trophoblasts in tissue sections. Human CCC and VCT at 5 gestational weeks also expressed HAVCR1, ITGA6, ITGA2, and S100P, as well as HSD3B and PTGES (Fig. 5G).

These data indicate that nCT and trophoblasts in the early post-implantation of cynomolgus monkey embryos and the first trimester placenta of human embryos share the same gene expression patterns, but pBAP do not.

### Primed PSC-derived cells express the genes related to amnion

Primate post-implantation EPI of embryo differentiates into the amnion in addition to the three germ layers (Ma et al., 2019; Xiang et al., 2020). It was also reported that primed PSCs corresponding to post-implantation EPI differentiate into amnion-like cells with BMP treatment (Zheng et al., 2019), hence, we assumed that primed PSCs might have the amnion-like property with BAP treatment. Ma et al. succeeded in culturing cynomolgus monkey embryos until gastrulation (Ma et al., 2019) and reported differentially expressed genes (DEGs) of the cynomolgus monkey amnion. We compared the gene expression patterns of DEGs of the amnion with pBAP, nTE, and nCT. Scatter plots show the trophoblast genes GATA2, GATA3, TFAP2C, TACSTD2, ENPEP, and DAB2 were commonly expressed in pBAP, nTE, and nCT (Fig. 6A). However, pBAP cells expressed over 200 amnion-specific genes of early post-implantation at significantly higher levels than nTE and nCT, showing that pBAP cells expressed amniotic genes more (Fig. 6A). TNC, CDH10, GABRP, IGFBP5, VTCN1, VIM, and HEY1 were identified as genes expressed at significantly higher levels in pBAP than in nTE or nCT (Fig. 6A). Next, we compared nTE, nCT, and pBAP to *in vivo* human amnion at more developed stage (9 gestational weeks) (Roost et al., 2015). A heat map revealed that amnion-predominantly expressed genes were highly upregulated in pBAP compared with nTE and nCT (Fig. 6B).

**Fig. 6.**
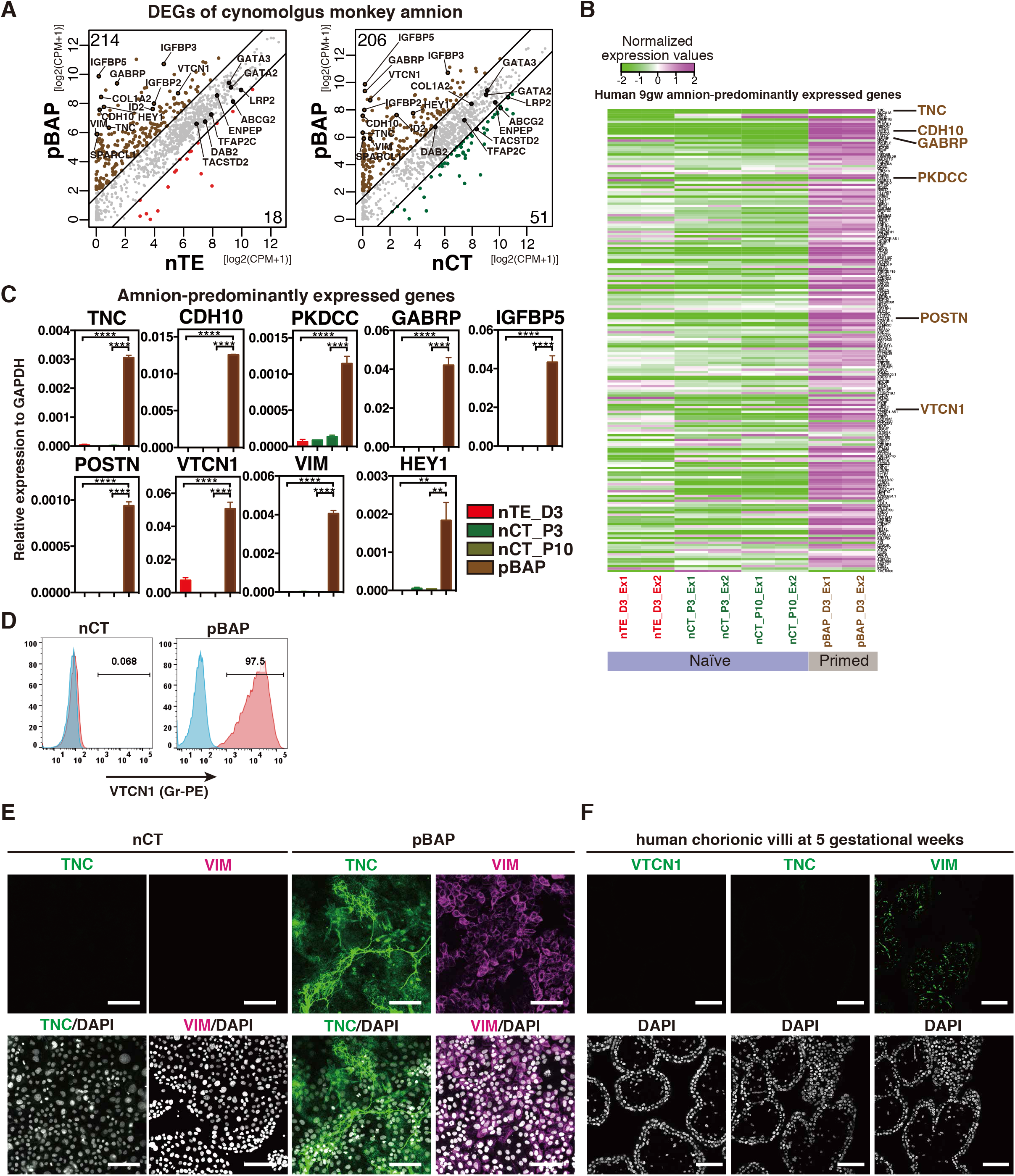
Primed PSC-derived cells express amnion-related genes. (A) Scatter plots of the average gene-expression levels for differentially expressed genes (DEGs) of cynomolgus monkey amnion between nTE (day 3) and pBAP (day 3) (left) and between nCT (passage 10) and pBAP (day 3) (right). The DEGs of amnion in cynomolgus monkey embryo cultures were defined previously (Ma et al., 2019). Brown circles indicate enriched genes in pBAP, red circles indicate enriched genes in nTE, and green circles indicate enriched genes in nCT. pBAP, nTE, and nCT share common genes including *GATA2, GATA3, TFAP2C, TACSTD2,* and *ENPEP.* Most of the enriched genes belong to pBAP (more than 200 genes). Solid black lines indicate 3-fold changes. Key genes are annotated, and the number of enriched genes are indicated in the top left of each plot. (B) Relative gene expression dynamics of nTE, nCT, and pBAP. Amnion-predominantly expressed genes were identified by comparing human amnion and placenta at 9 gestational weeks (Roost et al., 2015). The top 300 amnion-predominantly expressed genes and CPM > 1 in at least one sample are shown. Amnion-predominantly expressed genes were highly upregulated in pBAP compared to nTE and nCT. (C) RT-qPCR analysis of nTE, nCT, and pBAP for the key enriched genes identified in Figure 6A and 6B. n=3. (D) Flow cytometry analysis of nCT and pBAP for the pBAP-enriched gene VTCN1. Blue shows an unstained control. n=5. (E) Immunofluorescence images of nCT and pBAP (day 3) for the key pBAP-enriched genes shown in Figure 6A. TNC and VIM are stained in pBAP, but not in nCT. Scale bars, 100 μm. n=3. (F) Immunofluorescence images of human chorionic villi at 5 gestational weeks. Two key pBAP-enriched genes, VTCN1 and TNC, are not expressed, and VIM is stained in stromal cells but not in trophoblasts. Scale bars, 100 μm. n=3.

RT-qPCR confirmed that pBAP expressed the amnion-upregulated genes in Figure 6A and 6B much more strongly than nTE or nCT (Fig. 6C). Flow cytometry analysis showed that the cell surface antigen VTCN1 is expressed on pBAP but not on nCT (Fig. 6D), and immunofluorescence showed that pBAP expressed TNC and VIM, but nCT did not (Fig. 6E).

To examine whether the *in vitro* results agreed with *in vivo* development, we performed immunofluorescence staining of chorionic villi at 5 gestational weeks (Fig. 6F). VTCN1 and TNC were not detected in placenta. VIM was expressed in stromal cells but not in CCC, VCT, or ST. Based on the above, we concluded that pBAP bear similarity in gene expression to amniotic cells rather than CT.

### *In vitro* differentiation model captures trophoblast gene dynamics from pre-implantation to post-implantation

When the embryo implants into the uterus, TE contacts the endometrium and is thought to differentiate into CT, which migrates to the maternal decidua. We examined the dynamics of the gene expression using our model from the stages of nTE to nCT. To map the developmental coordinate axis of nTE, nCT, and previously reported TS cells (Cinkornpumin et al., 2020; Dong et al., 2020; Okae et al., 2018), we compared to cynomolgus monkey embryos during the peri-implantation period *in vivo* (Nakamura et al., 2016). The correlation coefficients for cynomolgus monkey embryos revealed that nTE_D2 have the highest correlation with late TE in the pre-implantation stage (PreL-TE) (Fig. 7A). As differentiation progressed, the correlation gradually shifted from pre-implantation to postimplantation (Fig. 7A). Reported naïve PSC-derived TS cells (Cinkornpumin et al., 2020; Dong et al., 2020) also had strong correlation to post-CT similar as nCT and CT30, suggesting that all naïve-, blastocyst-, and placenta-derived human TS cells are post-CT counterparts.

**Fig. 7.**
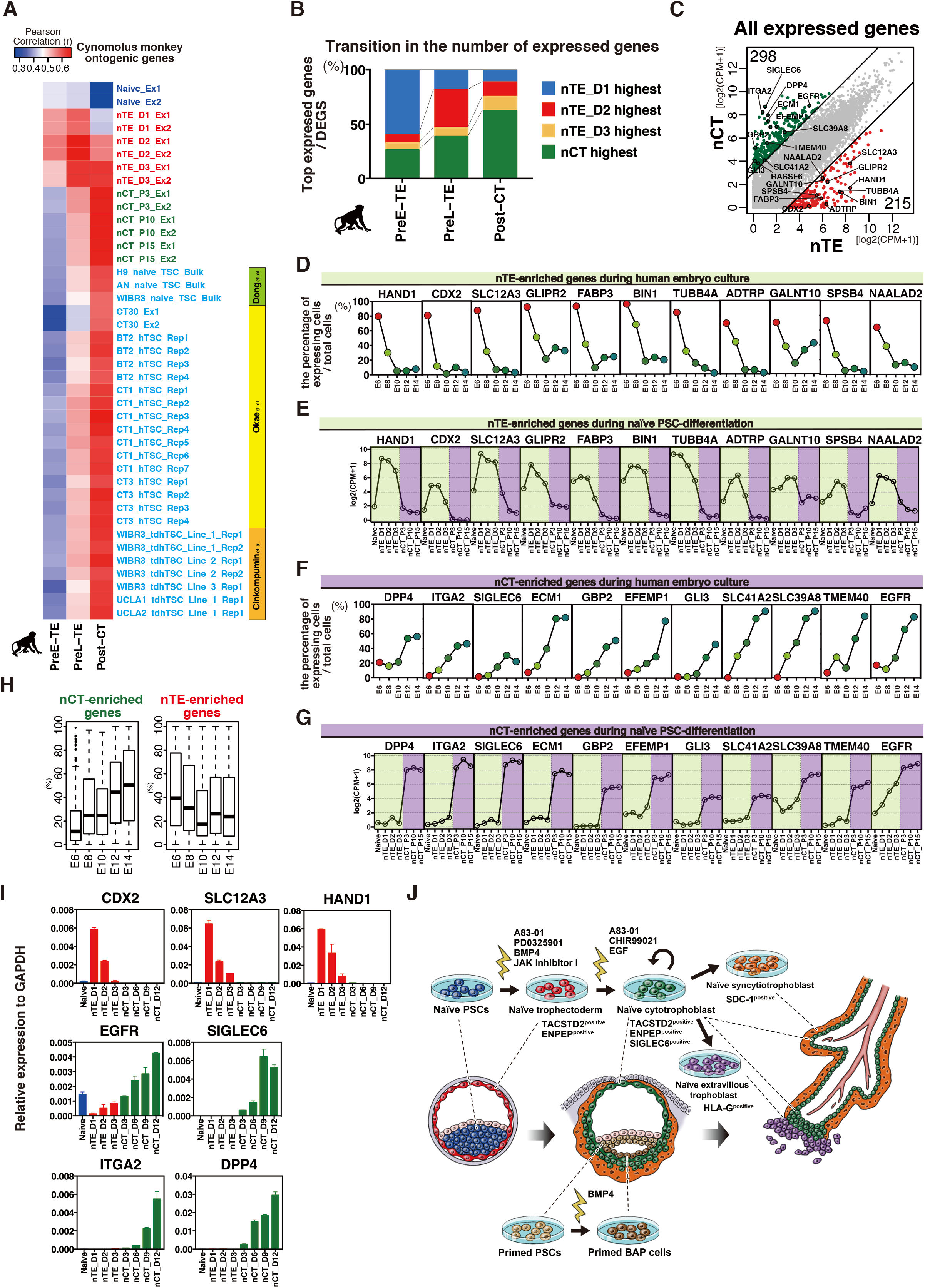
*In vitro* differentiation model captures trophoblast development from pre-implantation to post-implantation. (A) Heat maps of the correlation coefficients of naïve PSCs, nTE (days (D)1, 2, and 3), nCT (passages (p)3, 10 and 15), placenta-derived TS cells (Okae et al. 2018), blastocyst-derived TS cells (Okae et al. 2018), and reported naïve PSC-derived TS cells (Dong et al. 2020, Cinkornpumin et al. 2020) with cynomolgus monkey embryos. Analysis was performed using the cynomolgus ontogenic genes (Nakamura et al. 2016). RNA-seq data of trophoblast cells from Nakamura et al. was used for the analysis. PreE-TE, pre-implantation early trophectoderm; PreL-TE, pre-implantation late trophectoderm; Post-CT: post-implantation CT. CT1, CT3, and CT30; placenta-derived TS cells. BT2; blastocyst-derived TS cells. (B) The percentage of the top expressed differentially expressed genes (DEGs) during trophoblast development in nTE and nCT. The expression of each DEG in cynomolgus monkey trophoblast (PreE-TE, PreL-TE, Post-CT) was compared between nTE and nCT (passage 3). The distribution of nTE and nCT that expressed the DEGs most are shown. (C) Scatter plot of the average level for all expressed genes between nTE (day 2) and nCT (passage 3). Red circles indicate enriched genes in nTE, and green circles indicate enriched genes in nCT. Solid black lines indicate 8-fold changes. Key genes are annotated, and the number of enriched genes are indicated. (D) Key nTE-enriched gene expression dynamics during human embryo cultures. Human embryos were cultured from embryonic day (E)6 to E14 and were previously analysed by scRNA-seq (Zhou et al., 2019). The expression of the key nTE-enriched genes in Figure 7C was analysed in the trophoblast cells of human embryo cultures. The y-axis shows the percentage of expressing cells per total cells on each embryonic day. For example, on E6, around 80% of trophoblasts expressed *HAND1,* but almost no cells expressed *HAND1* at E10 and onward. The percentage of cells that expressed nTE-enriched genes decreased with human embryo development. (E) Key nTE-enriched gene expression dynamics during trophoblast differentiation from naïve PSCs. The expression peaked on days 1, 2 or 3 and then declined. The expression patterns are similar to those in Figure 7D. (F) Key nCT-enriched gene expression dynamics in human embryo cultures. Human embryos were cultured from E6 to E14 and analysed by scRNA-seq (Zhou et al., 2019). The expression of the key nCT-enriched genes in Figure 7C was analysed in the trophoblast cells of human embryo cultures. The y-axis shows the percentage of expressing cells per total cells on each embryonic day. The percentage of cells that expressed nCT-enriched genes increased with human embryo development. (G) Key nCT-enriched gene expression dynamics during trophoblast differentiation from naïve PSCs. The expression of key nCT-enriched genes was up-regulated during development. The expression patterns are similar to those in Figure 7F. (H) The expression dynamics of all nCT- and nTE-enriched genes (298 and 215 genes, respectively) during human embryo cultures. Human embryos were cultured from E6 to E14 and analysed by scRNA-seq (Zhou et al., 2019). The expression of all nCT- and nTE-enriched genes in Figure 6C was analysed in the trophoblast cells of human embryo cultures. The y-axis shows the percentage of expressing cells per total cells on each embryonic day. All enriched genes were combined. (I) The expression dynamics of genes enriched from nTE to nCT. TACSTD2^+^ENPEP^+^ (nTE_D3) were sorted on day 3 and re-cultured in ACE medium. *CDX2, SLC12A3* and *HAND1* expression was sharply decreased in ACE medium, whereas *EGFR, SIGLEC6, ITGA2* and *DPP4* expression was increased. (J) *In vitro* developmental model for trophoblast and trophoblast stem cells. After an embryo implant into the uterus, TE is thought to differentiate into CT, ST, and EVT. Naïve PSCs differentiate into TACSTD2^+^ENPEP^+^ nTE in A83-01, PD0325901, BMP4, and JAK inhibitor *in vitro.* nTE then differentiate into TACSTD2^+^ENPEP^+^SIGLEC6^+^ nCT in ACE medium. nCT can be maintained as trophoblast stem cells for more than 30 passages and differentiate into both HLA-G^+^ EVT and SDC1^+^ ST. The gene expression patterns indicate nCT correspond to *in vivo* proliferating CCC and VCT. Primed PSCs in BMP4 differentiate into pBAP, which express amnion-related genes.

We analysed the expression of DEGs of cynomolgus monkey pre-implantation early TE (PreE-TE), pre-implantation late TE (PreL-TE), and post-implantation CT (Post-CT) in nTE (D1, D2, D3) and nCT. Fig. 7B shows the percentage of the top gene expression among nTE and nCT: nTE_D1 expressed PreE-TE genes most, nTE_D2 expressed PreL-TE genes most, nCT expressed Post-CT genes dominantly, suggesting that nTE corresponds to pre-implantation TE, and nCT expressed early post-implantation trophoblast genes most (Fig. 7B).

Further, we compared gene expression between nTE_D2 and nCT_P3, looking at differences between pre-implantation TE and post-implantation CT (Fig. 7C). 215 genes were highly expressed in nTE, and 298 genes in nCT. The enrichment of Gene Ontology (GO) indicated that nTE-up-regulated genes are associated with development, while nCT-up-regulated genes are related to movement and migration (Fig. S7). These findings suggest that nCT expressed genes related to movement and migration as *in vivo* CT migrates to maternal decidua.

Next, we applied nTE- and nCT-enriched genes to human trophoblast development. Using the scRNA-seq data from E6 to E14 embryos (Zhou et al., 2019), we evaluated the expression of nTE- and nCT-enriched genes in the trophoblasts of cultured human embryos. First, we analysed he dynamics of 22 individual genes. The proportion of trophoblasts expressing nTE-enriched genes decreased during embryo culture (Fig. 7D). The gene expression dynamics from nTE to nCT showed similar down-regulation as cultured human embryos (Fig. 7E). nCT-enriched genes were also alike (Fig. 7F, 7G). The proportion of trophoblasts expressing nCT-enriched genes increased during embryo culture (Fig. 7F). nCT-enriched genes were upregulated from nTE to nCT, similar as cultured human embryos (Fig. 7G). Then we analysed the proportion of trophoblasts expressing nCT - or nTE-enriched genes (all of 298 and 215 genes). Trophoblasts that expressed nCT-enriched genes increased from E6 to E14 and trophoblasts that expressed nTE-enriched genes decreased, suggesting that our *in vitro* trophoblast model from nTE shows similar gene expression patterns as peri-implantation trophoblasts in human embryos (Fig. 7H).

Furthermore, we performed RT-qPCR to evaluate the precise gene expression dynamics from nTE to nCT of key enriched genes including CDX2, SLC12A3, and HAND1 for nTE-enriched genes and EGFR, SIGLEC6, ITGA2, and DPP4 for nCT-enriched genes (Fig. 7E). HAND1 and CDX2 were expressed in nTE and quickly decreased with development before the upregulation of SIGLEC6, while key enriched genes of nCT gradually increased in ACE medium.

## Discussion

We succeeded in establishing an *in vitro* model for the differentiation of naïve human PSCs into trophoblast lineage from TE to ST and EVT via CT (Fig. 7F). This model allows for observation of the entire trophoblast development from pre-implantation to post-implantation. To evaluate our model of the *in vitro* TE to CT transition, we compared it with *in vivo* cynomolgus monkey just after implantation and *in vitro* human and cynomolgus monkey cultured embryos as peri-implantation trophoblast. We also compared it with human first trimester placenta as a more developed stage of *in vivo* trophoblast, finding that our nTE-derived trophoblasts have high similarity with *in vivo* trophoblast development. Furthermore, we found that the gene expression patterns changed dramatically from pre-implantation TE to post-implantation CT. Our data also suggest the location of *in vivo* human TS cells in chorionic villi.

Naïve PSCs are thought to represent PSCs captured *in vitro* from pre-implantation EPI (Nichols and Smith, 2009). Although naïve mouse PSCs have existed for decades, naïve human PSCs were first reported only several years ago (Takashima et al., 2014; Theunissen et al., 2014). In this study, we show that naïve PSCs have functional distinctions compared with primed PSCs. We hypothesize that this functional difference is due to the developmental stage of the two PSCs. Naïve human PSCs have the capacity for differentiation into trophoblast lineage, whereas primed PSCs do not. Previously reported RNA-seq data indicated that the TE and EPI segregation of human embryos is later compared with mice (Petropoulos et al., 2016). Additionally, it is reported that TE in blastocysts could contribute to ICM *in vitro* (De Paepe et al., 2013), suggesting that it may be reasonable for naïve human PSCs to have TE differentiation potential.

At the peri-implantation stage in primate embryos, the amnion emerges from the embryonic disk (Ma et al., 2019; Niu et al., 2019; Xiang et al., 2020). Interestingly, previous reports suggested that the amnion expresses hCG, which is known to be expressed in ST (Plouzek et al., 1993; Thiede and Choate, 1963; Thiede and Fierer, 1966; Xiang et al., 2020). In addition, HLA-G, a major EVT marker, is also expressed in human amnion (Hammer et al., 1997; Strom and Gramignoli, 2016). These facts may explain why pBAP express hCG and HLA-G. Our re-analysis of previously reported RNA-seq data of cynomolgus monkey and human embryos revealed that trophoblast and amnion share many common genes. Indeed, we found many reported CT-specific genes are expressed in pBAP, including GATA3, TP63, and VGLL1 (Fig. 3D and S6E). However, our analysis revealed that genes such as IGFBP5 and GABRP in pBAP and HAVCR1 and ITGA2 in nCT were differentially expressed between amnion and trophoblast. Thus, our results show that cells induced from primed PSCs have more similar gene expression patterns to amnion than CT at post-implantation and first trimester (Fig. 7F). At the same time, several CT genes (HAVCR1, ITGA2, and SIGLEC6) are new and expected useful CT markers in addition to previously established trophoblast criteria (Lee et al., 2016).

The placenta of humans and cynomolgus monkeys have common features of chorionic villi, which distinguish them from the placenta of mice, which forms labyrinth structures. Therefore, we compared naïve PSC-derived trophoblasts to cynomolgus monkey trophoblasts. However, there are key species differences between humans and cynomolgus monkeys, including different gene expression patterns (Nakamura et al., 2016). Nevertheless, human research has many restrictions. To complement human data, it is essential to study animal models such as non-human primates that are close to humans, but also mice, which are generally advantageous as experimental animals.

Recently, several groups established culture models of human embryos beyond the implantation stage (Deglincerti et al., 2016; Shahbazi et al., 2016; Xiang et al., 2020; Zhou et al., 2019). However, culturing human embryos beyond 14 days is prohibited (Hyun et al., 2020; Sawai et al., 2020). Therefore, it is impossible to observe trophoblast development until terminal differentiation, and it is difficult to perform functional experiments. In the present study, we succeeded in purifying TE induced from naïve PSCs. We could therefore establish a culture model to reproduce the periimplantation TE-to-CT transition. Our results demonstrated that the gene expression of TE in preimplantation blastocyst is markedly different from those of CT in post-implantation embryos. This finding indicated that human nCT, recent naïve PSC-derived TS cells (Cinkornpumin et al., 2020; Dong et al., 2020), blastocyst- or placenta-derived TS cells (Okae et al., 2018), and trophoblast organoids (Haider et al., 2018; Turco et al., 2018) correspond to CT of post-implantation rather than pre-implantation TE. Furthermore, the comparison of gene expression patterns between nCT and chorionic villi revealed some cells of CCC and VCT are counterparts to CT stem cells. This finding is supported by previous reports that showed ITGA2^+^ cells in chorionic villi are proliferating trophoblasts that can differentiate into EVT and ST (Lee et al., 2018). Interestingly, our immunostaining revealed that the number of Ki67^+^ cells during the first trimester gradually decreased first in VCT and then in CCC. Finally, there are almost no Ki67^+^ cells in chorionic villi at 11 gestational weeks. This observation may suggest that the number of human TS or CT stem cell progenitors decreases during the first trimester and explain why human TS or CT stem cells can be established only from early first trimester placentas.

To conclude, our model allows us to analyse the TE-to-CT transition, which cannot be done using conventional TS cells. By using human PSCs including iPSCs, we can readily collect a large number of samples and perform various experiments including gene modifications. Our study thus contributes a major research tool to the study of human early development that is in agreement with current ethical regulations and extends research aimed at treatment including diseases related to the placenta.

## Acknowledgements

The authors are grateful to all patients for donating placental tissue for the research. The authors thank Naoyuki Kawamura, Akeo Kawamura (Kawamura Ladies Clinic), and Hironori Hamada (Adachi Clinic) for obtaining and documenting informed consent. The authors thank Dr. Hiroaki Okae and Dr. Takahiro Arima for kindly providing the CT30 cell lines and Dr. Keisuke Okita for kindly providing the 409B2 iPS cell line. The authors thank all Takashima Lab members for helpful discussions and assistance. We also thank Dr. Kanae Mitsunaga for the flow cytometry analysis, Mr. Shunsuke Kihara for the imaging analysis, and Tim Smith, Maria G Martin (University of Cambridge), and Ai Hirase-Hirabayashi (Kyoto University) for technical support. The authors are grateful to Dr. Peter Karagiannis for critical reading and English editing of the manuscript. This work was supported by MEXT KAKENHI (Grant Number:16H02465) and AMED (Grant Number: 19bm0704035, 19bm0104001) to Y.T., and Grant-in-Aid for JSPS Research Fellow (Grant Number:JP19J13503) to S.I.

## Author Contributions

S.I. and Y.T. conceived the study, designed the experiments, and interpreted the data. S.I. performed the majority of the experiments. K. S., M.K, T.Y., and T.N performed the RNA-seq analysis. Y.I., K. S., and B.K performed the cell cultures and molecular work. C.I., I.O., Y.K., T.N., H.T, and M.S. performed the cynomolgus monkey experiments. N.M performed the electron microscopy. The manuscript was written by S.I. and Y.T. M.S, T.Y, E.K., and M.M. edited the manuscript.

---

**Fig. S1.**
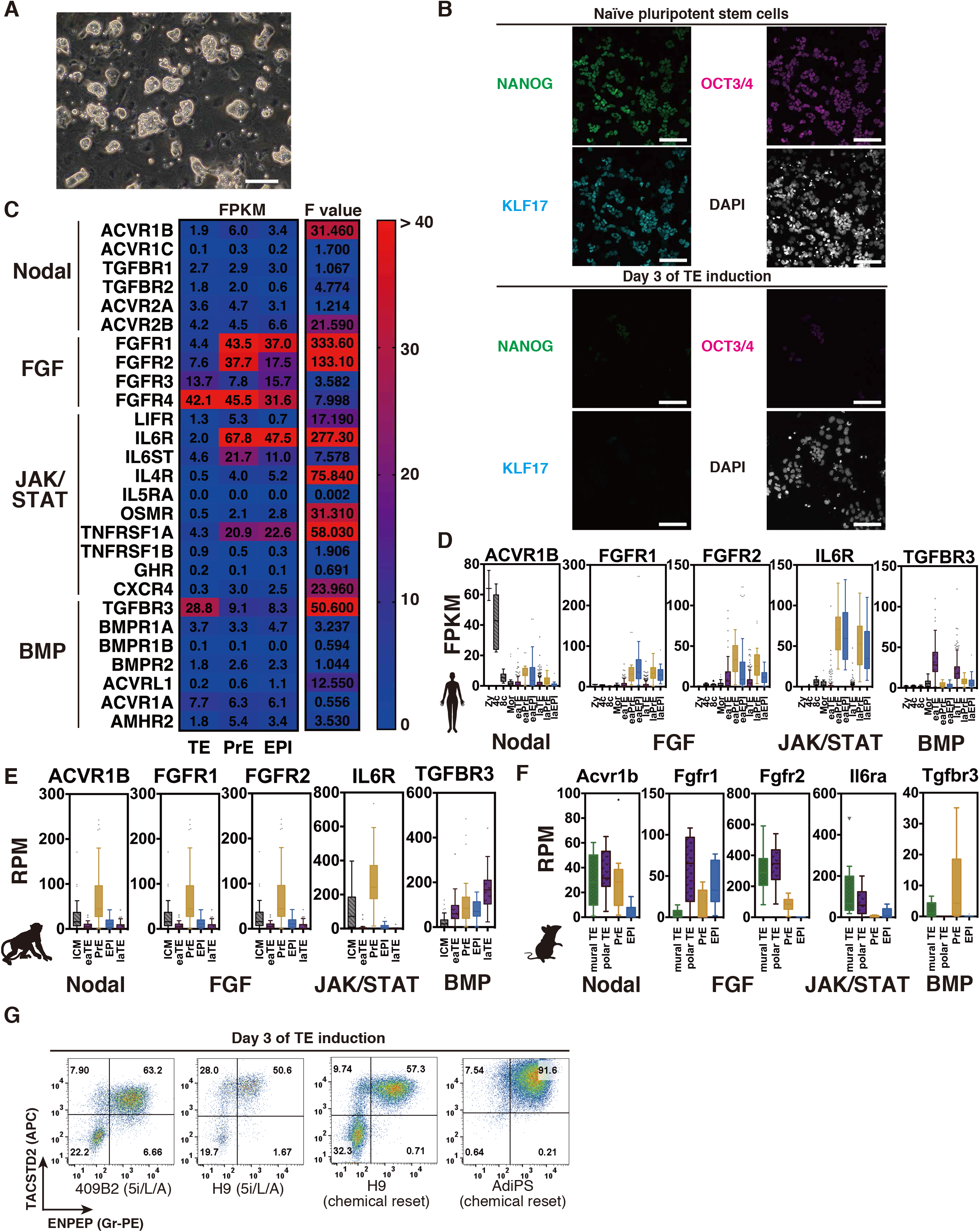
Generation of trophectoderm-like cells using chemical compounds. Related Figure 1. (A) Morphology of naïve human PSCs. H9 PSCs were cultured in t2iLGö. Scale bar, 100 μm. Representative images from n=10. (B) Immunofluorescence images of pluripotency markers. Naïve PSCs and naïve PSC-derived cells on day 3 after the trophectoderm (TE) induction were stained by NANOG, OCT3/4, and KLF17. White: DAPI. Scale bars, 100 μm. Representative images from n=3. (C) Gene expression of receptors and F values. Representative receptors for the Nodal, FGF, JAK/STAT, and BMP signaling pathways in human blastocysts. The heat maps show the gene expression (FPKM; fragments per kilobase of transcript per million mapped reads) of samples from TE, primitive endoderm (PrE) and epiblast (EPI). F values were calculated by one-way analysis of variance. (D) Gene expression in human pre-implantation embryo (embryonic days (E)1-7). Box and whiskers plots in the style of Tukey showing FPKM values of the representative receptors for Nodal, FGF, JAK/STAT, and BMP signaling. Receptors with the highest F values among each signaling group are shown. Zy, zygote; 4c, 4 cells; 8c, 8 cells; Mor, morula; eaTE, early TE (E5); eaPrE, early primitive endoderm ((PrE), E5); eaEPI, early epiblast ((EPI), E5); laTE, late TE (E7); laPrE, late PrE (E7); laEPI, late EPI (E7). Previously reported single-cell RNA-sequencing (scRNA-seq) data was used for the analysis (Stirparo et al., 2018). (E) Gene expression in cynomolgus monkey pre-implantation embryo. Box and whiskers plots in the style of Tukey showing RPM (reads per million-mapped reads) values of representative receptor expression for Nodal, FGF, JAK/STAT, and BMP signaling. ICM, inner cell mass; eaTE, early trophectoderm; PrE, primitive endoderm; EPI, epiblast; laTE, late trophectoderm. These lineage stages were annotated by Nakamura et al. (Nakamura et al., 2016) (GSE74767). (F) Gene expression in mouse pre-implantation embryo (E4.5). Box and whiskers plots in the style of Tukey showing RPM values of representative receptor expression for Nodal, FGF, JAK/STAT, and BMP signaling. mural TE, mural trophectoderm; polar TE, polar trophectoderm; PrE, primitive endoderm; EPI, epiblast. These lineage stages were annotated by Nakamura et al. (Nakamura et al., 2015) (GSE63266). (G) Flow cytometry analysis for the expression of TACSTD2 and ENPEP on day 3 after the TE induction of naïve human PSCs (409B2 iPSCs reset by 5i/L/A; H9 ESCs reset by 5i/L/A; H9 reset by chemical reset; and AdiPS reset by chemical reset). Both TACSTD2 and ENPEP are expressed in all cell lines on day 3. Representative images from n=3.

**Fig. S2.**
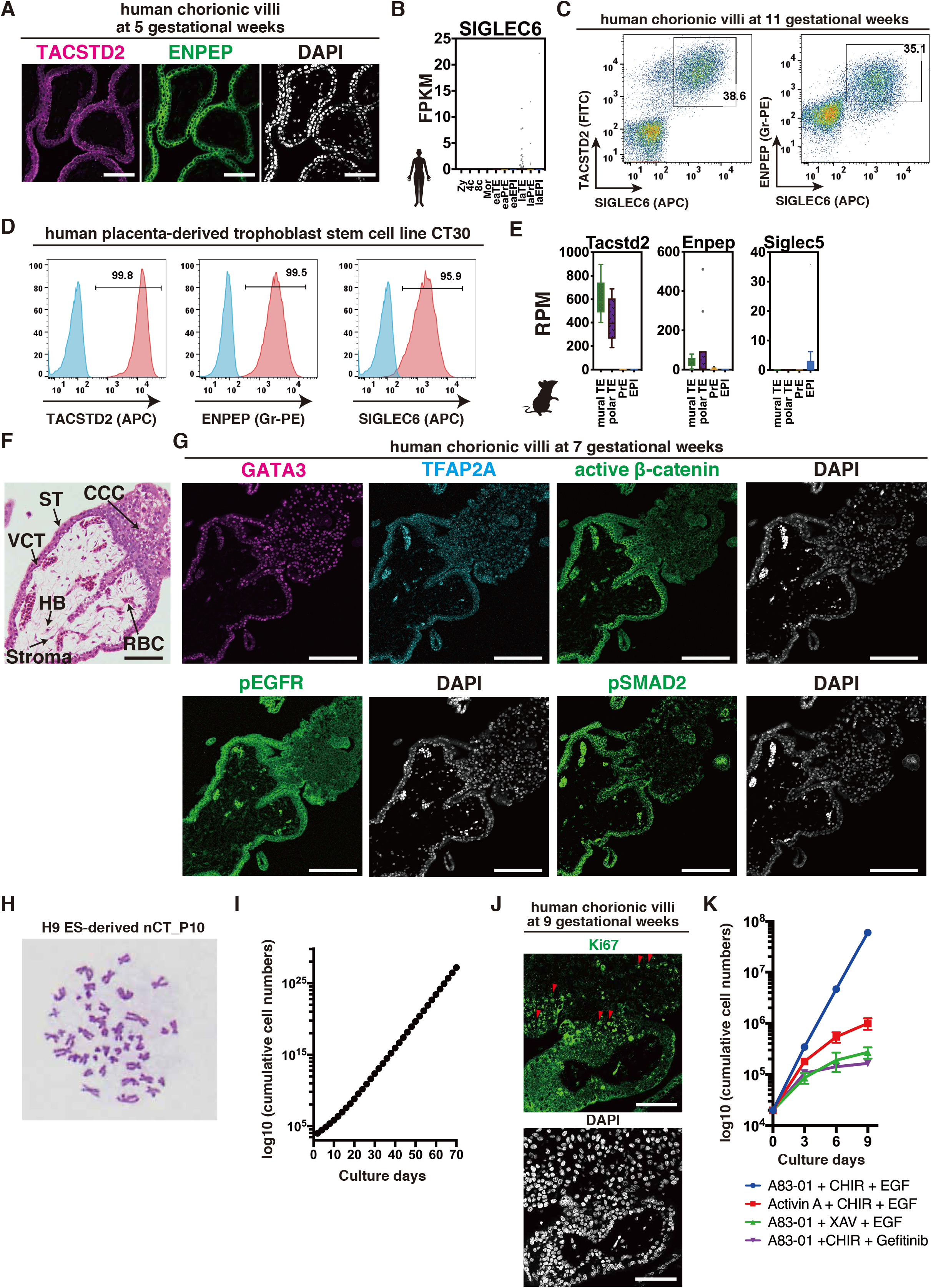
nCT can be maintained as CT-like stem cells. Related Figure 2. (A) Immunofluorescence images of human chorionic villi at 5 gestational weeks for TACSTD2 and ENPEP. Scale bars, 100 μm. Representative images from n=3. (B) Expression in FPKM values of SIGLEC6 (CD327) for human early embryo (E1-7). SIGLEC6 is not expressed in human pre-implantation embryo. Zy, zygote; 4c, 4 cells; 8c, 8 cells; Mor, morula; eaTE, early TE (E5); eaPrE, early primitive endoderm ((PrE), E5); eaEPI, early epiblast ((EPI), E5); laTE, late TE (E7); laPrE, late PrE (E7); laEPI, late EPI (E7). Previously reported single-cell RNA-sequencing (scRNA-seq) data was used for the analysis (Stirparo et al., 2018). (C) Flow cytometry analysis for the expression of TACSTD2, ENPEP, and SIGLEC6 on human chorionic villi (11 gestational weeks). (D) Flow cytometry analysis for the expression of TACSTD2, ENPEP, and SIGLEC6 on the human placenta-derived trophoblast stem cell line CT30 (Okae et al., 2018). Blue shows an unstained control. Representative images from n=3. (E) Expression in RPM values of Tacstd2, Enpep, Siglec5 (mouse ortholog for human SIGLEC6) in mouse pre-implantation embryo. mural TE, mural trophectoderm; polar TE, polar trophectoderm; PrE, primitive endoderm; EPI, epiblast. These lineage stages were annotated by Nakamura et al. (Nakamura et al., 2015) (GSE63266). (F) Hematoxylin-Eosin staining of human chorionic villi at 7 gestational weeks. CCC, cytotrophoblast cell column; VCT, villous cytotrophoblast; ST, syncytiotrophoblast; HB, Hofbauer cell (villous macrophage); RBC, fetal nucleated red blood cell. Scale bars, 100 μm. Representative images from n=5. (G) Immunofluorescence images of human chorionic villi at 7 gestational weeks. GATA3, TFAP2A, active β-catenin, phosphorylated EGF receptor, and phosphorylated SMAD2 are shown. Active β-catenin was localized in the cell membrane of CCC and VCT. Phosphorylated EGF receptor was strongly expressed in the cell membrane of CCC, VCT and ST. Phosphorylated SMAD2 was not accumulated in the nuclei of CCC, VCT, or ST. Scale bars, 100 μm. Representative images from n=3. (H) A representative result for the Giemsa staining of H9 naïve human ES cell-derived CT stem cells at passage 10 bearing normal karyotype (46, XX). 30 cells were analysed. (I) Proliferation of nCT. nCT in ACE medium was passaged every 2 days. nCT continued to proliferate as CT-like stem cells. (J) Immunofluorescence images of chorionic villi at 9 gestational weeks for the proliferation marker Ki67. Ki67 was expressed in the nuclei of some parts of CCC and VCT. Scale bars, 100 μm. Representative images from n=3. (K) Three compounds (A83-01, CHIR99021, and EGF) are essential to maintain nCT. nCT were cultured in three compounds including one opposing factor of ACE. Each opposing factor, Activin A, XAV (Wnt/β-catenin signaling inhibitor), and Gefitinib (EGFR tyrosine kinase inhibitor), inhibits the proliferation of nCT. nCT was cultured in N2B27 supplemented with some combination of 1 μM A83-01, 2 μM CHIR99021, 50 ng/ml EGF, 50 ng/ml Activin A, 2 μM XAV939, and 2 μM Gefitinib on iMatrix 511 silk. CHIR, CHIR99021; XAV, XAV939.

**Fig. S3.**
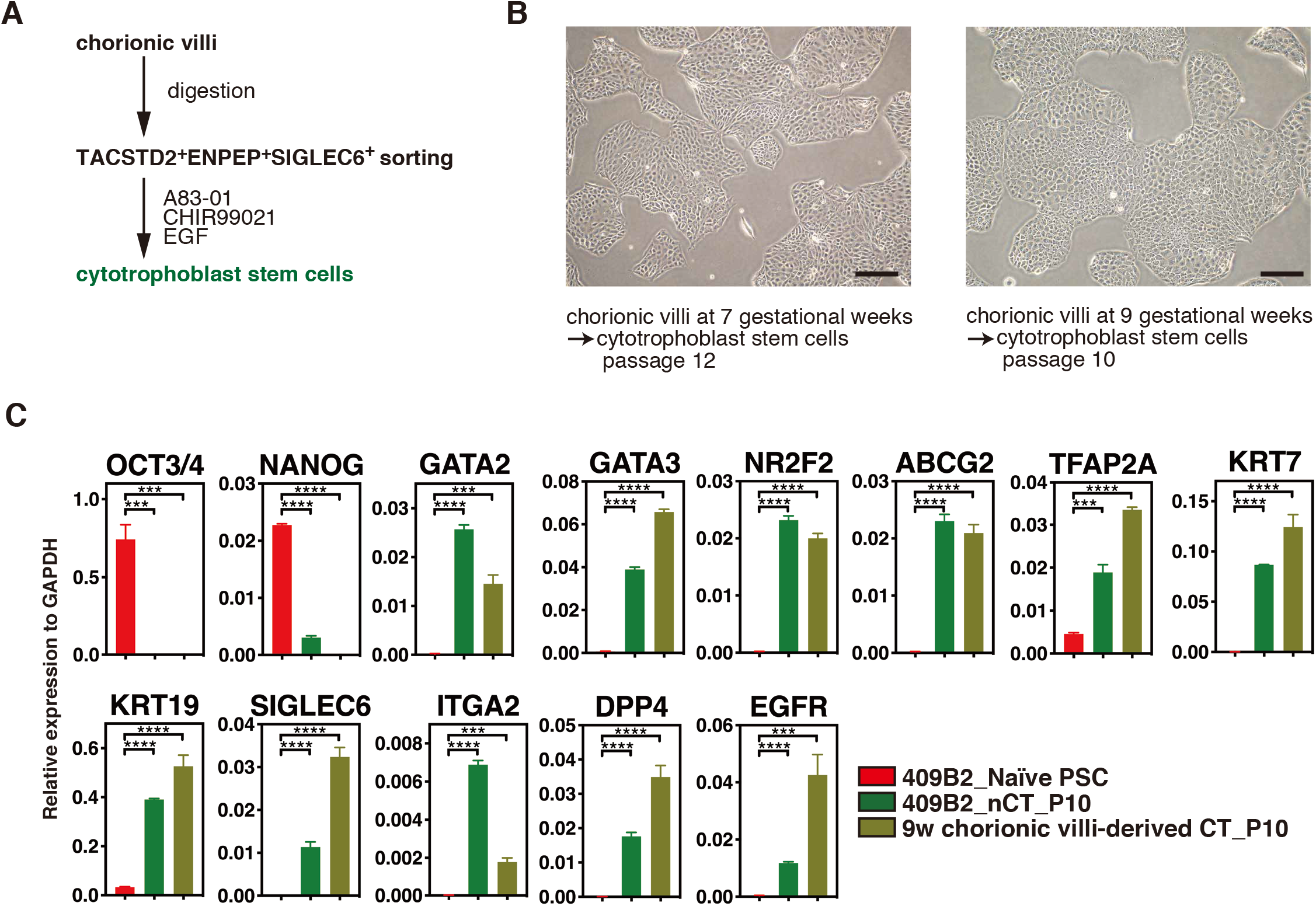
The derivation of new TS cell lines from chorionic villi in three compounds (A83-01, CHIR99021, and EGF). Related Figure 2. (A) Schema of the derivation of the TS cell line from chorionic villi. TACSTD2^+^ENPEP^+^SIGLEC6^+^ cells were sorted from chorionic villi at 7 and 9 gestational weeks and cultured in ACE condition. (B) The morphology of TS cells in ACE at passage 12 (7 gestational weeks) and passage 10 (9 gestational weeks). Representative images from n=3. (C) RT-qPCR analysis for CT markers in TS cells. nCT and TS cells at passage 10 were analysed. 409B2; 409B2 human iPS cell line. n=3.

**Fig. S4.**
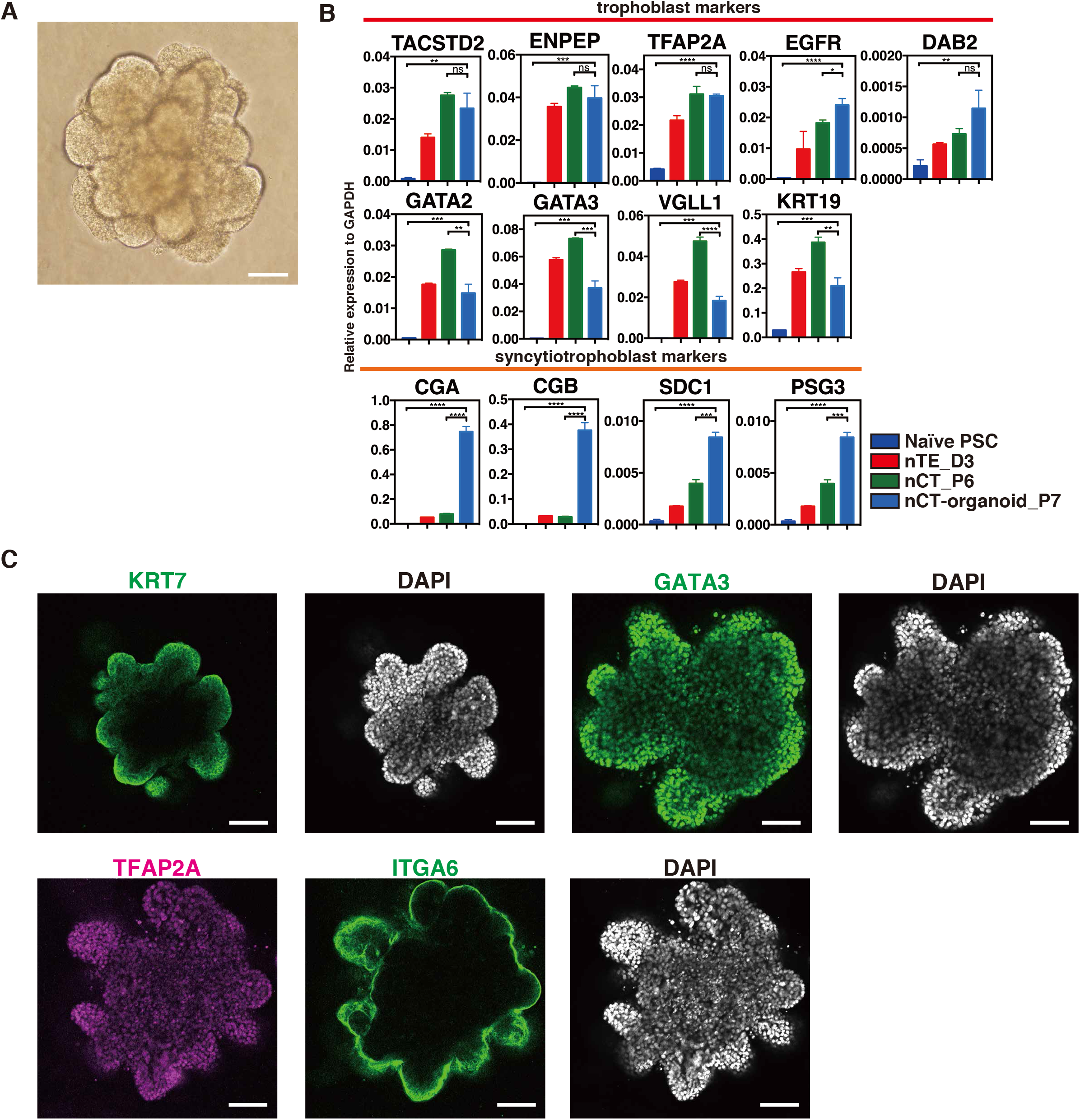
nCT can be maintained as nCT-organoids. Related Figure 2. (A) Morphology of a three-dimensional trophoblast organoid. nCT was cultured in trophoblast organoid medium (Turco et al., 2018). Scale bar, 100 μm. Representative images from n=10. (B) RT-qPCR analysis of CT and ST markers in trophoblast organoids. nCT passage 6 and nCT-organoid passage 7 were analysed. n=3. (C) Immunofluorescence images of nCT-organoid for the trophoblast markers KRT7, GATA3, TFAP2A, and ITGA6. Scale bars, 100 μm. Representative images from n=3.

**Fig. S5.**
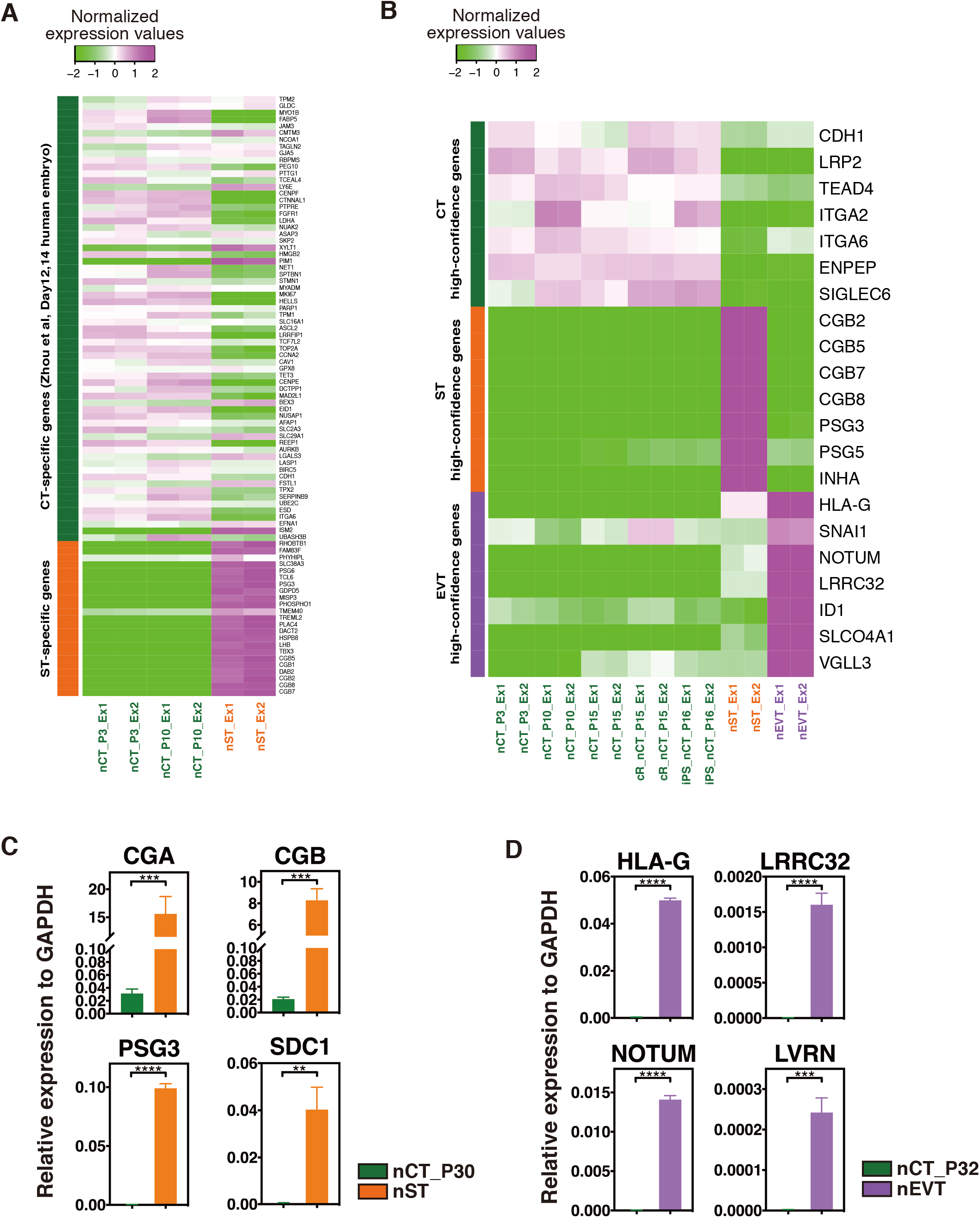
nTE-derived CT differentiates into ST and EVT. Related Figure 3. (A) Relative gene expression dynamics of nCT and nST. Specific genes for CT and ST on E12 and E14 were determined by Zhou et al. (Zhou et al., 2019). Human embryos cultured *in vitro* were analysed using single-cell RNA-sequencing (scRNA-seq) (Zhou et al., 2019). (B) Relative gene expression dynamics of high-confidence stage markers in nCT, nST, and nEVT. We assigned 7 genes to each stage by the expression and distribution in scRNA-seq data of the placenta. CT-, ST-, EVT-representative genes were confirmed by the maternal-fetal interface (https://maternal-fetal-interface.cellgeni.sanger.ac.uk/) (Vento-Tormo et al., 2018). (C) RT-qPCR analysis of *CGA*, *CGB, PSG3,* and *SDC1.* After 30 passages maintained in ACE medium, nCT were cultured in Forskolin. ST marker genes were upregulated after 6 days of induction. n=3. (D) RT-qPCR analysis of *HLA-G, LRRC32, NOTUM,* and *LVRN.* After 32 passages maintained in ACE medium, nCT were cultured in A83-01, Neuregulin-1, and Geltrex. EVT marker genes were upregulated after 8 days of induction. n=3.

**Fig. S6.**
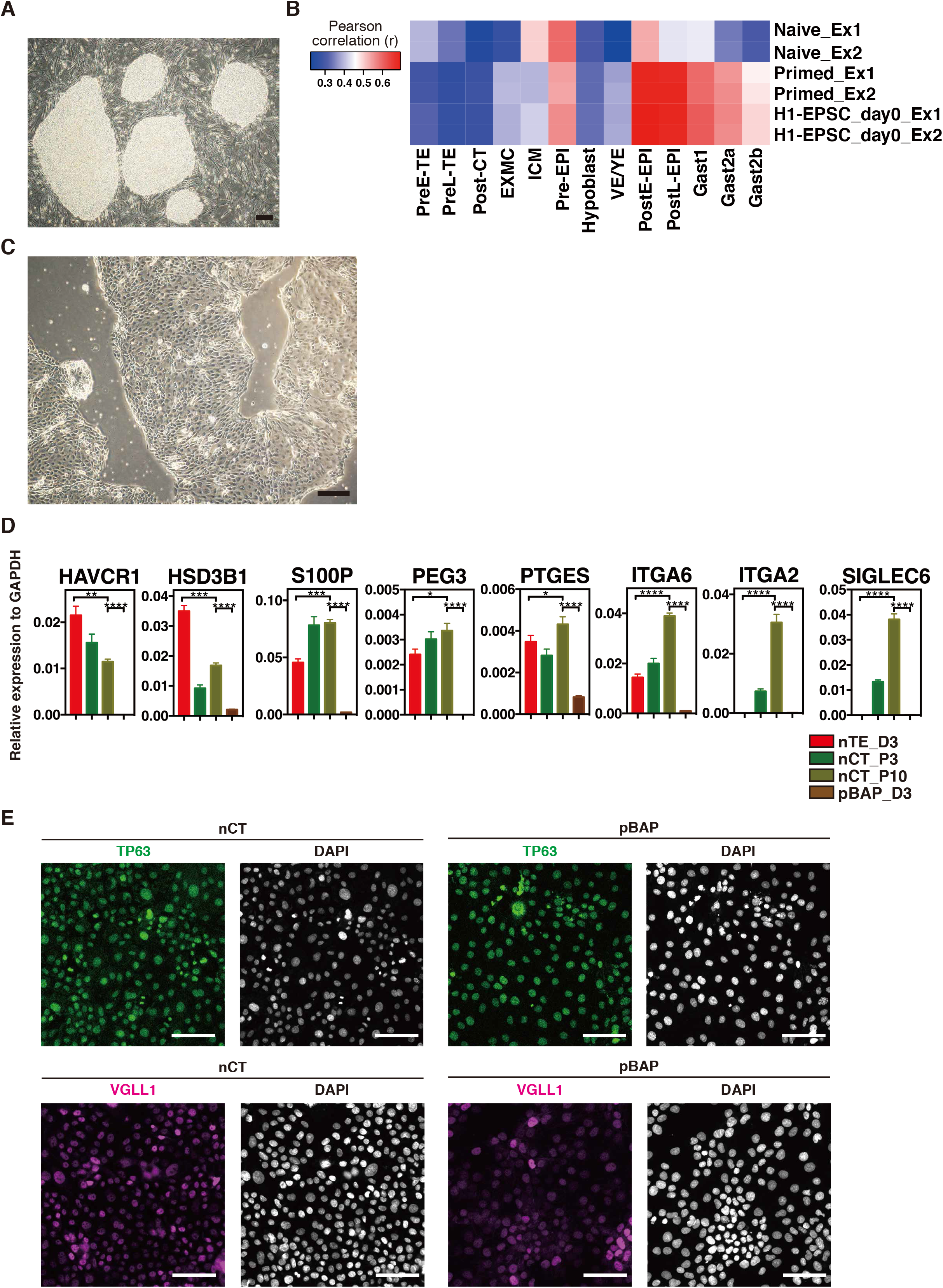
nCT represents post-implantation CT of *in vivo* humans and cynomolgus monkeys. Related Figure 5. (A) Morphology of primed human PSCs. H9 primed PSCs were cultured in KSR/bFGF. Scale bar, 100 μm. Representative images from n=10. (B) Heat maps of the correlation coefficients among naïve PSCs, primed PSCs, and EPSCs on day 0 with cynomolgus monkey embryos. Primed PSCs and EPSCs have a strong correlation with post-implantation early or late EPI (PostE-EPI and PostL-EPI). RNA-seq data from Nakamura et al. (Nakamura et al., 2016) was used for the analysis. Nakamura et al. classified cells from cynomolgus embryos into pre-implantation early or late TE (PreE-TE and PreL-TE, respectively), postimplantation CT (Post-CT), ICM, visceral endoderm/yolk sac endoderm (VE/YE), pre-implantation EPI (Pre-EPI), post-implantation early or late EPI (PostE-EPI and PostL-EPI, respectively), or gastrulating cells (early: Gast1, middle: Gast2a, and late: Gast2b). (C) Morphology of pBAP cultured in ACE medium for 10 passages. Scale bar, 100 μm. Representative images from n=3. (D) mRNA expression of nTE, nCT and pBAP for key trophoblast-enriched genes. RT-qPCR analysis of nTE, nCT, and pBAP for the key trophoblast-enriched genes identified in Figure 5B. (n=3) (E) Immunofluorescence images of TP63 and VGLL1 in nCT and pBAP. TP63 and VGLL1 are known as trophoblast markers. However, nCT and pBAP express TP63 and VGLL1 similarly. Scale bars, 100 μm. (n=3)

**Fig. S7.**
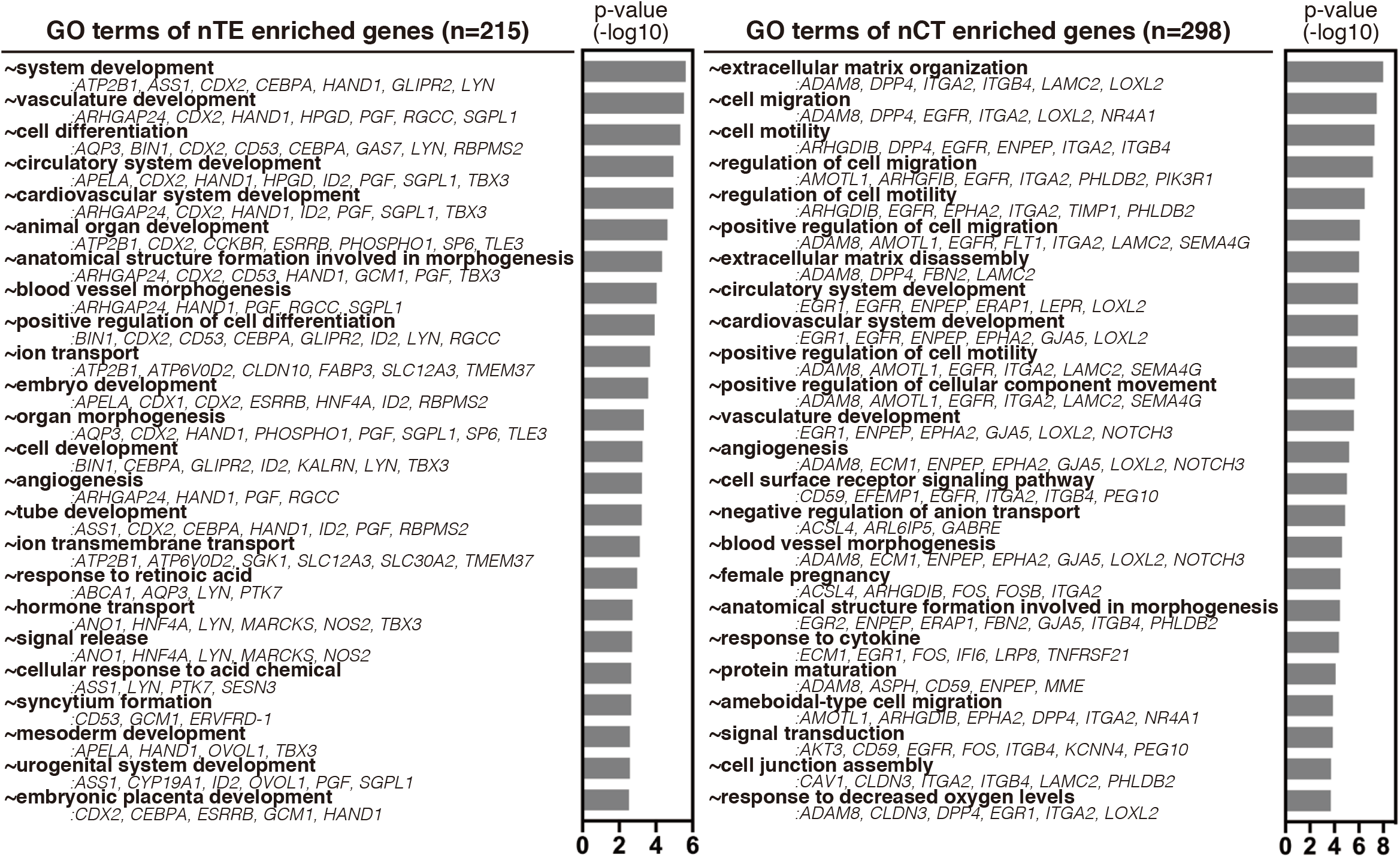
The character of nTE- and nCT-enriched genes. Related Figure 7. Enrichments of GO terms and their representative genes in nTE- and nCT-enriched genes are indicated.

## Methods

### Experimental models and Subject Details

All cells were cultured in 5% O_2_, 5% CO_2_ except where noted otherwise.

### Pluripotent Stem Cell Culture

#### Culture of human primed pluripotent stem cells

The human ESC line H9 (WiCell Research Institute), human iPSC lines 409B2 (Okita et al., 2011) and AdiPS (Takashima et al., 2014) were cultured on mouse embryonic fibroblast (MEF) cells (1×10^6^ cells per 6-well plate).

Primed human PSCs were maintained in DMEM/Ham’s F-12 (Nacalai Tesque, Cat. 08460-95) with 20% Knockout Serum Replacement (KSR; Thermo Fisher Scientific, Cat. 10828028), 1% nonessential amino acids (Thermo Fisher Scientific, Cat. 11140-050), 4 ng/ml recombinant human basic fibroblast growth factor 2 (bFGF; Oriental Yeast, Cat. NIB 47079000), and 0.1 mM 2-mercaptoethanol (2ME; Sigma-Aldrich, Cat.M3148). Cultures were passaged every 5-7 days as small clumps by dissociation buffer containing 0.025% Trypsin (Thermo Fisher Scientific, Cat. 15090-046), 1 mg/ml Collagenase IV (Thermo Fisher Scientific, Cat. 17104-019), 20% KSR, and 1 μM CaCl2.

Naïve human PSCs were maintained in t2iLGö medium, which consists of N2B27 (Ndiff227: Takara Bio, Cat. Y40002) with 1 μM PD0325901 (PD03; Tocris, Cat. 4192), 1 μM CHIR99021 (CH; Sigma-Aldrich, Cat. SML1046), 10 ng/ml Recombinant human LIF (hLIF; Peprotech, Cat. 300-05), and 2 μM Gö6983 (Gö; Tocris, Cat. 2285), as previously described (Takashima et al., 2014). Naïve PSCs were passaged every 3-5 days using Accutase (Sigma-Aldrich, Cat. A6964).

Chemical conversion to naïve PSCs was performed as previously described (Guo et al., 2017). Primed PSCs (1×10^4^ cells/cm^2^) were seeded on MEF feeder cells in primed PSC medium with 10 μM Y-27632. On the next day, the medium was switched to cRM-1 (N2B27 + 1 μM PD03, 10 ng/ml hLIF, and 1 mM valproic acid sodium salt (VPA; Sigma-Aldrich, Cat. P4543)). On day 3, the medium was replaced with cRM-2 (N2B27 + 1 μM PD03, 10 ng/ml hLIF, 2 μM Gö, and 2 μM XAV939 (Sigma-Aldrich, Cat. X3004)). Dome-shaped naïve colonies were observed around two weeks after plating. Cells were split every 5–7 days after dissociation with Accutase. Reset cells were passaged and maintained on MEF feeders in t2iLGö. Chemical conversion to naïve PSCs using 5iLA was also performed as previously described (Theunissen et al., 2014, 2016). 2×10^5^ cells/cm^2^ were seeded on MEF feeder cells under primed PSC medium with 10 μM Y-27632. On the next day, the medium was switched to 5iLA media (Ndiff227 supplemented with 1 μM PD03, 1 μM CH, 1 μM WH-4-023 (A Chemtek, Cat. H620061), 0.5 μM SB590885 (R&D systems, Cat. 2650), 10 μM Y-27632, 10 ng/ml hLIF, and 20 ng/ml Activin A (R&D, Cat. 338-AC-010)). After conversion to naïve PSCs, the cells were maintained in t2iLGö on MEF feeder cells.

### Cynomolgus Monkey Specimens

Experimental procedures using cynomolgus monkeys were approved by the Animal Care and Use Committee of Shiga University of Medical Science. The procedures in cynomolgus monkeys for oocyte collection, intra-cytoplasmic sperm injection (ICSI), pre-implantation embryo culture, and transfer of pre-implantation embryos into foster mothers were performed as described previously with minor modifications (Nakamura et al., 2016; Yamasaki et al., 2011). Ovulation induction with follicle stimulating hormone (Gonapure; ASKA Pharmaceutical) was performed by embedding an implantable and programmable micro-fusion device (iPRECIO; Primetech Corporation) subcutaneously. The day when the ICSI was performed was designated as embryonic day (E) 0. The progression of the pre-implantation development of cynomolgus monkeys was highly similar to that of rhesus monkeys (Wolf et al., 2004), but was somewhat slower than that of humans (Wong et al., 2010). For embryo transfer, 4 to 5 two-cell to blastocyst-stage embryos were selected and transferred into an appropriate recipient female. For the detection of pregnancy of early post-implantation embryos, implanted embryos were monitored by ultrasonography around E13 and the implanted uterus was surgically removed and bisected for the isolation of embryos.

### Clinical Samples

All tissue samples used for this study were obtained with written informed consent from all participants in accordance with the guidelines in The Declaration of Helsinki 2000. Human chorionic villi of elective abortions were obtained from Kawamura Ladies Clinic and Adachi Hospital (5-11 gestational weeks). The tissue donations were made entirely voluntarily by women undergoing termination of pregnancy. Donors were asked to give explicit written consent for the fetal material to be collected, and only after they had been counselled about the termination of their pregnancy. Gestational age was estimated with a combination of last menstrual period and ultrasonography. The use of all human prenatal samples for the work described in this study was approved by ethics committee in Center for iPS Cell Research & Application (CiRA), Kyoto University.

### Methods details

### Differentiation into nTE

Naïve PSCs were seeded on Laminin-E8 (0.15 μg/cm^2^ iMatrix 511 silk; Nippi) at 2 x 10^4^ cells/cm^2^. The Laminin-E8-coated dish was pre-incubated at 37 °C for at least 30 min. The initial induction medium was composed of Ndiff227, 2 μM A83-01 (ALK4/5/7 inhibitor; Tocris), and 2 μM PD03. The following day, the medium was changed to Ndiff277, 2 μM A83-01, 2 μM PD03, and 1 μg/ml JAK inhibitor I (Merck, Cat. 420099-1 MGCN). The medium was changed again the next day. 10 ng/ml recombinant human BMP4 (R&D systems) was optionally added for 24 hours after the induction. Cells were dissociated with Accutase for 20-30 min.

### Culture of nCT

TACSTD2^+^ENPEP^+^ nTE cells on day 3 were sorted and cultured in ACE medium (Ndiff227, 1 μM A83-01, 2 μM CH, and 50 ng/ml EGF (R&D systems, Cat.236-EG)) on Laminin-E8 (0.15 μg/cm^2^) at 4 x 10^4^ cells/cm^2^). The culture medium was replaced every two days. Cells were passaged every three to six days by dissociation with Accutase for 10-15 min and seeded at a 1:3-1:4 split ratio. 10 μM Y-27632 was added for every passage. Cells were cryopreserved in Stem-Cellbanker (Takara Bio) and stored in a deep freezer at −80 °C.

### Differentiation into nST

nCT was seeded on Col IV (1 μg/ml) or Laminin-E8 (0.15 μg/cm^2^) at 1 x 10^4^ cells/cm^2^. ST medium was composed of DMEM/Ham’s F-12 supplemented with 0.1 mM 2ME, 0.3% BSA, 1% ITS-X supplement (Life Technologies), 4% KSR, 2 μM forskolin (cAMP activator; Wako, Cat. 067-02191), and 2.5 μM Y-27632 as previously described (Okae et al., 2018). ST medium was replaced at day 3 and cultured for three additional days. Cells were incubated in 21% O2, 5% CO_2_ in a humidified incubator at 37 °C.

### Differentiation into nEVT

nCT was seeded on Col IV (1 μg/ml) or Laminin-E8 (0.15 μg/cm^2^) at 0.8 x 10^4^ cells/cm^2^. EVT-1 medium was composed of DMEM/Ham’s F-12 supplemented with 0.1 mM 2ME, 0.3% BSA, 1% ITS-X supplement, 4% KSR, 7.5 μM A83-01, 100 ng/ml NRG1 (Cell Signaling, Cat. 5218SC), and 2.5 μM Y-27632 as previously described (Okae et al., 2018). After the cells were plated, Geltrex (Thermo Fisher Scientific, Cat. A1413302) was added to a final concentration of 2%. EVT-1 medium was replaced on day 3 with EVT-2 medium (DMEM/Ham’s F-12 supplemented with 100 mM 2ME, 0.3% BSA, 1% ITS-X supplement, 4% KSR, 7.5 μM A83-01, and 2.5 μM Y-27632), and Geltrex was added to a final concentration of 0.5%. On day 6 after the EVT induction, the cells were dissociated into single cells with Accutase for 10-15 min and seeded on a new Col IV- or Laminin-E8-coated plate at a 1:2-1:3 split ratio. The cells were cultured in EVT-3 medium (DMEM/Ham’s F-12 supplemented with 100 mM 2ME, 0.3% BSA, 1% ITS-X supplement, 7.5 μM A83-01, and 2.5 μM Y-27632), and Geltrex was added to a final concentration of 0.5%. The cells were analysed on day 8 after the EVT induction and incubated in 21% O2, 5% CO_2_ in a humidified incubator at 37 °C.

### Culture of nCT-organoids

nCT-organoids were cultured as trophoblast organoids (Turco et al., 2018). 2-4 x 10^5^ nCT cells were resuspended in 500 μL growth-factor-reduced Matrigel (Corning, Cat. 356231) on ice. Matrigel drops (25 μl) including nCT cells were plated per well into a 48-well culture plate (Cellstar, Cat. 677-180), which was then incubated in a humidified incubator at 37 °C for 15 min. The firm drops were overlaid with trophoblast organoid medium (Ndiff227 supplemented with 1.25 mM N-Acetyl-L-cysteine (Sigma, Cat. A2921201), 2 mM L-glutamine, 0.5 μM A83-01, 1.5 μM CH, 50 ng/ml EGF, 80 ng/ml R-spondin 1 (R&D systems, Cat. 4645-RS-100), 100 ng/ml bFGF, 50 ng/ml HGF (Peprotech, Cat. 100-39), 2.5 μM PGE2 (Sigma, Cat. P0409), and 2.5 μM Y-27632). Trophoblast organoid medium was replaced every 2-3 days. Organoids were incubated in 5% O2, 5% CO_2_ in a humidified incubator at 37 °C and passaged every 7-10 days by mechanical disruption with a Picus NxT electronic single channel electronic pipette (Sartorius) on a mix cycle of 99 rounds (4-5 times), maximum speed. The organoids were cryopreserved in Stem-Cellbanker (Takara Bio) and stored in a deep freezer at −80 °C.

### Culture of BAP-treated primed pluripotent stem cells

Human primed PSCs were dissociated into single cells with trypsin/EDTA (Nacalai Tesque). The cells were plated on a Laminin-E8-coated dish at a density of 2 x 10^4^ cells/cm^2^ and cultured with MEF-conditioned medium supplemented with 10 ng/ml BMP4, 1 μM A83-01, and 0.1 μM PD173074 (FGF2 signaling inhibitor; Selleck, Cat. S1264), as described previously with some modifications (Amita et al., 2013). DMEM/Ham’s F-12 medium containing 0.1 mM 2ME, 1% ITS-X supplement, 1% NEAA, 2 mM L-glutamine, and 20% KSR was cultured on MEF for 24 hours, and the supernatant was collected and used as MEF-conditioned medium. The medium was changed daily.

### Culture of human trophoblast stem cells

CT30 cells were cultured as described previously (Okae et al., 2018). The culture medium was composed of DMEM/Ham’s F-12 supplemented with 0.1 mM 2ME, 0.3% BSA, 0.2% FBS, 1% ITS-X supplement, 1.5 μg L-ascorbic acid (Sigma-Aldrich), 0.5 μM A83-01, 2 μM CH, 50 ng/ml EGF, 1 μM SB431542 (Tocris), 0.8 mM VPA, and 5 μM Y-27632. The cells were cultured on a Col IV-coated well dish and passaged every 2-3 days by dissociation with TrypLE at a 1:2-1:4 split ratio. Cells were incubated in 21% O2, 5% CO_2_ in a humidified incubator at 37 °C. Cells were cryopreserved in Cell Banker 1 (Takara Bio) and stored in a deep freezer at −80 °C.

### Culture of JAR

JAR cells were cultured as described previously. The cells were plated on gelatin-coated dish at a density of 5 x 10^3^ cells/cm^2^ and cultured with RPMI-1640 medium supplemented with 10% FBS. The medium was changed every 2 days. The cells were incubated in 21% O2, 5% CO_2_ in a humidified incubator at 37 °C. At subconfluency, cells were passaged by dissociation with trypsin/EDTA for 5-15 min and seeded at a 1:3-1:6 split ratio. The cells were plated, cryopreserved in Cell Banker 1 (Takara Bio) and stored in a deep freezer at −80 °C.

### Isolation of human cytotrophoblast

Whole chorionic villi were cut into small pieces and enzymatically digested three times in a solution containing 0.25% Trypsin-EDTA solution, 1 mg/ml collagenase IV, 200 U/mL DNase (Sigma-Aldrich), 25 mM HEPES, and DMEM/F’12 medium with agitation at 37°C. Pooled cell suspensions were filtered through a 70 μm mesh filter (Corning). After the lysis of erythrocytes in RBC lysis solution (Santa Cruz Biotechnology), CT was purified using a Alexa Fluor 488-conjugated anti-TACSTD2 antibody, a PE-conjugated anti-CD249 antibody, and biotin-conjugated anti-SIGLEC6 antibody. The TACSTD2^+^ENPEP^+^SIGLEC6^+^ fraction was collected for RNA-seq samples of in vivo CT.

### Bisulfite sequencing of ELF5 promoter

Genomic DNA was isolated with the DNeasy Blood & Tissue kit (Qiagen), and 100-400 ng of genomic DNA was processed for bisulfite conversion using the EpiTect Bisulfite kit (Qiagen) following the manufacturer’s instructions. Nested PCR was performed with Quick Taq HS DyeMix (Toyobo) on a Veriti 96-Well Thermal Cycler (Applied Biosystems). The first PCR (10 cycles) was carried out with ELF5-2b (ELF5-201) BiS-483F and +31R primers. Ten percent of the first PCR products were used for the second PCR (35 cycles) at an annealing temperature of 50 °C using ELF5-2b BiS-432F and - 3R primers. The gel-purified PCR amplicons were inserted into TOPO-XL PCR cloning vectors (Thermo Scientific), and the products were used to transform Library Efficiency DH5_a_ Chemically Competent Cells (Thermo Scientific). After purification with ExoSAP-IT Express PCR Cleanup Reagents (Thermo Scientific), at least eight clones were sequenced for each sample.

### Immunocytochemistry

Cells and organoids were fixed with 4% paraformaldehyde (PFA, Nakalai Tesque) solution for 10 min at room temperature, permeabilized with 1% BSA/PBS + 0.5% Triton X-100 for 1 hour at room temperature, and blocked with 5% donkey serum (Jackson ImmunoResearch, Cat. 017-000-121)/PBS + 0.05% Tween-20 (PBS-BT) for 2 hours at room temperature. Primary antibodies were diluted in PBS-BT and incubated overnight at 4 °C. After two washes with PBS-BT, the cells were incubated in PBS-BT for 12 hours at 4 °C. Secondary antibodies were AlexaFluor conjugated, diluted 1:1000 in PBS-BT, and incubated overnight at 4 °C. Nuclei were stained with 4,6-diamidino-2-phenylindole (DAPI, Sigma-Aldrich). Images were acquired using a TSC-8 (Leica, Germany) or LSM710 (Zeiss, Germany) confocal laser scanning microscope.

### Immunohistochemistry

Cynomolgus monkey embryos in vivo and human clinical samples were fixed in 10% buffered formalin for 24 hours and embedded in paraffin. Serial sections were taken every 200 μm. After the sections were deparaffinized, microwave antigen retrieval was performed by incubation with Histo VT one (Nakalai Tesque) for 20 min. After tissue sections were permeabilized with 1% BSA/PBS + 0.5% Triton X-100 for 1 hour at room temperature, they were incubated with PBS-BT for 30 min at room temperature. Subsequently, primary antibodies were diluted in PBS-BT and incubated overnight at 4 °C. Secondary antibodies were diluted 1:500 in PBS-BT and incubated overnight at 4 °C. Nuclei were stained with 4,6-diamidino-2-phenylindole (DAPI) and mounted in VECTASHIELD mounting medium (Vector Laboratories). Images were acquired using a TSC-8 or LSM710 microscope.

### Conventional scanning electron microscopy

Monolayer cell cultures on small glass coverslips (ø 6mm) were chemically fixed with 2.5% glutaraldehyde (GA) and 2% PFA in NaHCa buffer (including 100 mM NaCl, 30 mM HEPES, 2 mM CaCl2, adjusted at pH 7.4 with NaOH) for at least a few hours at room temperature. After several washes with 0.1 M cacodylate buffer (pH7.4), the cultures were proceeded to post-fixation with 1% osmium tetroxide/1% potassium ferrocyanide in 0.1 M cacodylate buffer for 1 hour. After several washes in distilled water, a second post-fixation was performed with 5% uranyl acetate solution overnight at 4 °C in a dark refrigerator and then dehydrated with a series of ethanol washes (from 70% to absolute 100%). Finally, after special treatment with hexamethyldisilazane (HMDS, Sigma-Aldrich, St. Louis USA), the dried specimens were sputter-coated with pure gold (EMITECH k950K, Quorum Technologies, Kent UK). Fine observation was performed using a FEI Quanta FEG250 scanning electron microscope (Thermo Fischer Scientific, Oregon USA).

### Karyotype analysis

H9 naïve human ES cell-derived CT stem cells at passage 10 of culture with ACE medium were incubated with metaphase arrest solution (Genial Helix) for 1.5 hours. The treated cells were dissociated into single cells, resuspended by 0.075 M KCl prewarmed to 37 °C, and incubated for 10 minutes. After cell fixation using 75% methanol and 25% acetic acid, the fixed cells were dropped on the slides and blow-dried. The slides were stained in Giemsa solution. Karyotype images (30 cells were analysed) were obtained with a BX-51 microscope (OLYMPUS, Tokyo, JAPAN), and the number of chromosomes was counted manually.

### Flow cytometry

After dissociation by Accutase at 37 °C, the cells were incubated in basal medium for 30 min at 37 °C. After washes with HBSS (Thermo Fisher Scientific) and BSA (Sigma-Aldrich) (1% BSA/PBS), the cells were incubated with 1% BSA/PBS for 30 min on ice. Primary and secondary antibody incubations were performed for 30 min each in 1% BSA/PBS on ice, and the cells were washed one time with 1% BSA/PBS. Streptavidin-APC (Biolegend) was used as the secondary antibody for biotin-conjugated primary antibodies. Flow cytometry analysis and cell sorting was performed using a BD LSR Fortessa (BD Biosciences, United States) or a BD FACSAria II (BD Biosciences), and the data were analysed using FlowJo software (LLC).

### Quantitative real-time PCR (mRNA & miRNA)

Cellular RNA was extracted using the RNeasy Mini kit (Qiagen) following the manufacturer’s instructions. MiRNA was extracted using the Direct-zol RNA Miniprep (Zymo Research). First-strand cDNA was synthesized from 1000 ng RNA using SuperScript IV Reverse Transcriptase (Thermo Fisher Scientific) according to the manufacturer’s instructions. cDNA for miRNA quantitative real-time PCR was synthesized using the miScript II RT kit (Qiagen). The cDNA was diluted 20-fold. Real-time PCR reactions were performed with the Quantstudio 12K Flex real-time PCR system and Quantstudio 3 real-time PCR system (Applied Biosystems, Foster City, CA, USA) using PowerUp SYBR Green Master Mix (Applied Biosystems) for mRNA and the miScript SYBR Green PCR kit (Qiagen) for miRNA. Samples for mRNA were run as technical triplicates, and the results were analysed using the delta-delta cycle threshold approach with GAPDH as the endogenous control. In the same way, results for miRNA were analysed using the delta-delta cycle threshold approach with RNU6-2 as the endogenous control. All original primers for mRNA were designed using PrimerBlast (http://www.ncbi.nlm.nih.gov/tools/primer-blast/). TaqMan Assays (Applied Biosystems) for hsa-miRNA were performed in miR-518c-5p, hsa-miR-520c-3p, and hsa-miR-519d. The results were analyzed using QuantStudio Design&Analysis Software v1.4.1 (Thermo Fisher Scientific).

### Fusion event analysis

The fusion index represents the percentage of cell-cell fusion events. Syncytia were defined as cells with at least three nuclei. The number of DAPI-stained nuclei and syncytia were counted. The fusion index was calculated using the following formula: [(N-S)/T] x 100. N, the number of nuclei in the syncytia; S, the number of syncytia; T, total number of nuclei counted.

### ELISA of hCG

nCT cells were seeded on Laminin-E8 (0.15 μg/cm^2^) at a density of 1 x 10^4^ cells/cm^2^ and differentiated into nST as described above. ST medium was replaced at day 3. The supernatant was collected at day 6 and stored at −80 °C until use. As controls, nCT cells were seeded on Laminin-E8 (0.15 μg/cm^2^) at a density of 1 x 10^4^ cells/cm^2^ and cultured in ACE medium. On day 3, the supernatant was collected and stored at −80 °C until use. hCG ELISA (Abcam; Cat. ab100533) was performed using 100 μl supernatant in triplicate alongside hCG standards according to the manufacturer’s instructions. Absorbance at 450 nm and 570 nm was monitored by a multimode plate reader (2104 Envision; PerkinElmer Inc, Waltham, MA, USA). As a positive control, maternal serum hCG was also examined in the same experiment. The concentration of secreted hCG in the supernatants was calculated from the line formula of the standard plots in Graphpad Prism.

### Time-lapse analysis

Time-lapse phase-contrast microscopy was performed using Biostation CT (Nikon-Intech, Tokyo, JAPAN). Images were acquired every 40 minutes over a course. MP4 movies were generated using CL-Quant Ver3.31 (Nikon-Intech, Tokyo, JAPAN).

### RNA sequencing and data analysis

Total RNA was isolated with the miRNeasy Mini kit. The purity of the extracted RNA was evaluated using a NanoDrop 2000 spectrophotometer (Thermo Fisher Scientific), and the concentration of the RNA was assessed using a Qubit 3.0 fluorometer (Thermo Fisher Scientific). The library was constructed by using 200 ng total RNA and the TruSeq Stranded mRNA LT Sample Prep Kit (Illumina). Sequencing was performed on a NextSeq 500 using the High Output v2.5 kit (75 cycles, FC-404-2005; Illumina). The sequenced reads were trimmed to remove low-quality bases and adaptor sequences using cutadapt-1.16 (Martin, 2011). The trimmed reads were mapped to the human reference genome (hg38) using HISAT v2.1.0 with GENCODE v30 (Frankish et al., 2019). The uniquely mapped reads were summarized at the gene level using HTSeq-count v0.11.2 (Anders et al., 2015), and the expression level of each gene was normalized using edgeR v3.24.3 (McCarthy et al., 2012). Low-expression genes across all samples were excluded from subsequent analyses. Regarding to our RNAseq data, CPM (counts per million reads) values were used as gene expression levels, whereas FPKM values were used only for comparison analysis with cynomolgus monkeys (Figure 5A, 7A, S6). For heatmap, expression values were normalized to the median or mean of all data sets or a specific condition. Representations of heat maps, correlation analyses, hierarchical clustering analyses, and principal component analyses (PCA) were performed using R software (v3.5.1). Gene Ontology analysis was performed using DAVID 6.8 (Huang da et al., 2009).

### Comparison of gene expression between humans and cynomolgus monkeys

To compare the gene expression between humans and cynomolgus monkeys, common genes listed in the cynomolgus monkeys-humans one-to-one annotation table were used as described previously (Nakamura et al., 2016; Nakamura et al., 2017).

### Quantification and Statistical Analysis

Quantification methods are described in the figure legends or detailed here in STAR METHODS. The exact number of replicates used are indicated within individual figure legends. Results are expressed as the means, and error bars represent SD unless otherwise indicated. If not otherwise indicated, statistical significance was determined by unpaired two-tailed Student’s t test using GraphPad Prism 7 or R (****, p ≤ 0.0001; ***, p ≤ 0.001; **, p ≤ 0.01; *, p ≤ 0.05, ns = not significant). In all cases, the data was assumed to meet t test requirements (normal distribution and similar variance), and differences were considered to be statistically significant when P values were less than 0.05.

## Data and materials availability

The RNA sequencing data from this study are being deposited in NCBI Gene Expression Omnibus.

